# Functional γδT-omics pipeline reveals compartmentalization of Vδ1⁺ T cell migration, tumor-reactivity, and clonality in human colorectal cancer

**DOI:** 10.1101/2025.08.19.671055

**Authors:** Lucrezia C.D.E. Gatti, Mara J.T. Nicolasen, Farid Keramati, Peter Brazda, Angelo D. Meringa, Dorine A. De Bont, Tineke Aarts-Riemens, Willemijn J. C. Brandwijk, Maud I.M. van der Wijst, Anniek H.G. Stuut, Laia Gasull Celades, Daniel Zawal, Konstantinos Vazaios, Annet Daudeij, Maarten A. Huismans, Maria A. Parigiani, Trudy Straetemans, Dennis X. Beringer, Onno Kranenburg, Christopher Berlin, Susana Minguet, Rebecca Kesselring, Hendrik G. Stunnenberg, Jeanine M.L. Roodhart, Zsolt Sebestyen, Jürgen Kuball

**Affiliations:** Center for Translational Immunology, University Medical Center Utrecht, Utrecht University, The Netherlands; Princess Maxima Center for Pediatric Oncology, Utrecht, The Netherlands; Center for Molecular Medicine, University Medical Center Utrecht, Utrecht University, The Netherlands; Faculty of Biology, University of Freiburg, Freiburg, Germany; Signalling Research Centres BIOSS and CIBSS, University of Freiburg, Freiburg, Germany; Center of Chronic Immunodeficiency (CCI) and Institute for Immunodeficiency, University Clinics and Medical Faculty, Freiburg; Department of Hematology, University Medical Center Utrecht, The Netherlands; Laboratory of Translational Oncology, Division of Imaging and Cancer, University Medical Center Utrecht, The Netherlands; Department of Surgical Oncology, Division of Imaging and Cancer, University Medical Center Utrecht, The Netherlands; Utrecht Platform for Organoid Technology, Utrecht University, The Netherlands; Department of General and Visceral Surgery, Medical Center-University of Freiburg, Faculty of Medicine, University of Freiburg 79106 Freiburg im Breisgau, Germany; German Cancer Research Center (DKFZ), 69120 Heidelberg, Germany; German Cancer Consortium (DKTK) partner site Freiburg, 79106 Freiburg im Breisgau, Germany; Department of Molecular Biology, Faculty of Science, Radboud University, Nijmegen, The Netherlands; Department of Medical Oncology, University Medical Centre Utrecht, Utrecht University, Utrecht 3508 GA, The Netherlands

**Author notes:** First authors contributed equally (Lucrezia C.D.E. Gatti, Mara J.T. Nicolasen). Last authors contributed equally (Zsolt Sebestyen, Jürgen Kuball).

## Abstract

γδT cells play a pivotal role in cancer immune surveillance, yet the current knowledge of their function across the compartments in solid tumors is meager. To address this gap, we developed a comprehensive γδT-omics platform that integrates functional screening, biomimetic migration assays, γδTCR repertoire analysis, and transcriptomic profiling. Using matched samples from 31 patients with microsatellite-stable colorectal cancer (CRC), we analyzed γδT cells from peripheral blood (PBLs_γδ_), adjacent colon (LPLs_γδ_), primary tumors (pTILs_γδ_), and liver metastases (mTILs_γδ_). This approach uncovered striking compartmentalization of γδT cell phenotypes, clonality, and function. Tumor-reactive, clonally expanded Vδ1⁺ γδT cells were enriched in primary tumors and shared transcriptional and functional features with lamina propria lymphocytes (LPLs). In contrast, Vδ1⁺ γδT cells from liver metastases lacked tumor reactivity, exhibited distinct γδTCR repertoires, and expressed transcriptional signatures associated with TGF-β-mediated suppression and cellular quiescence, suggesting they are shaped by tissue-specific environmental cues. CXCL16 secretion by tumor cells initiated Vδ1⁺ LPLs_γδ_ migration, which was further amplified by γδTCR-mediated CCL5 induction from pTILs_γδ_, leading to CCR5 downregulation and subsequent entrapment of pTILs_γδ_ within the tumor microenvironment. Accordingly, our clinical data from an independent second cohort of 69 patients showed that infiltration by pTILs_γδ_, but not mTILs_γδ_, is associated with a protective effect against CRC progression. In summary, our study offers a compartment-resolved perspective on γδT cell behavior in CRC, revealing key trafficking and functional mechanisms, and enabling the identification of novel tumor-reactive γδTCRs and migratory cues to inform immunotherapeutic strategies for both primary and metastatic CRC.

**Graphical abstract:** 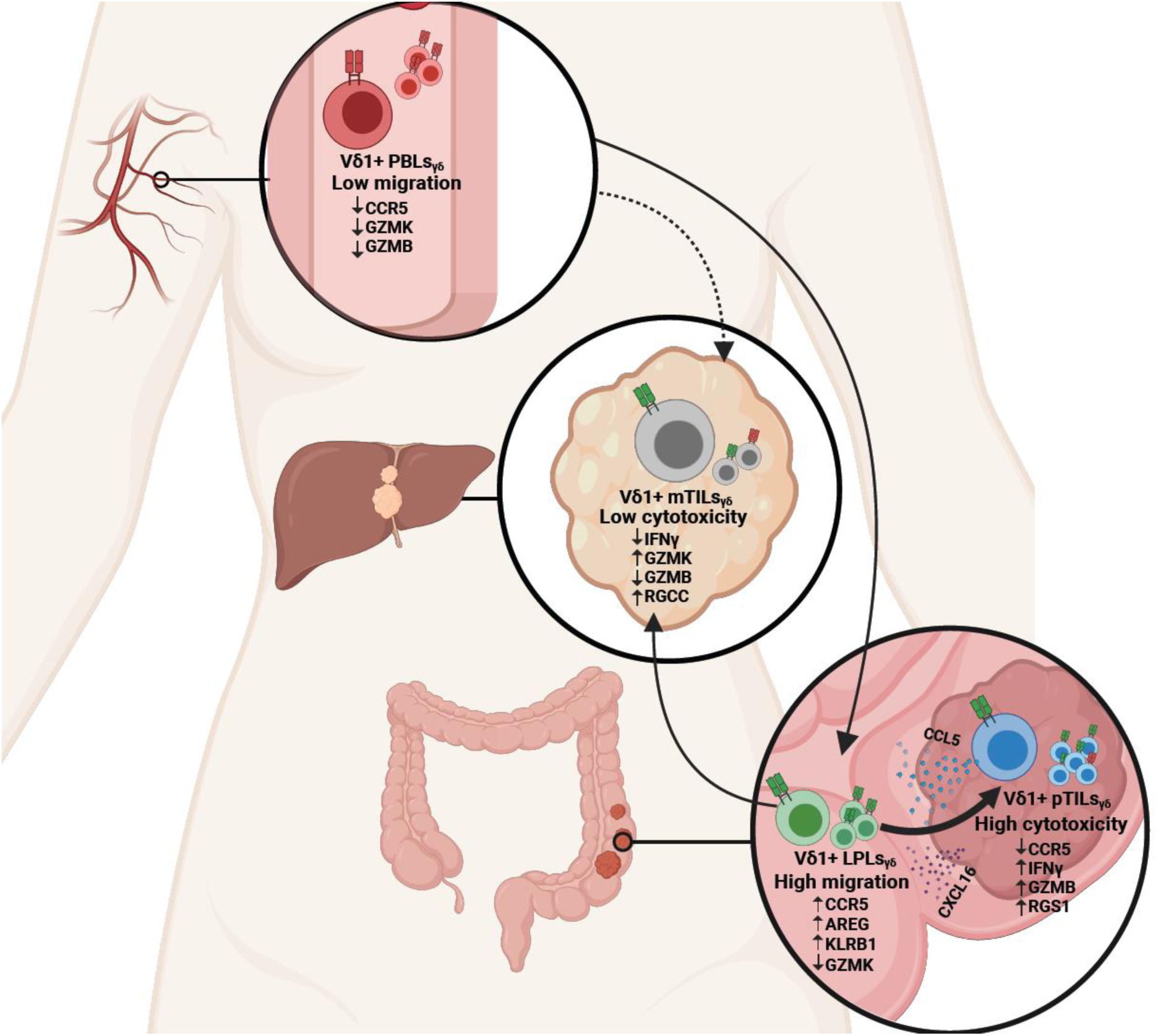

## Introduction

Colorectal cancer (CRC) is one of the leading causes of cancer-related mortality worldwide, with significant impacts on global health. While early detection and improved treatments have increased the survival rate, the overall mortality rate remains high, particularly for late-stage diagnoses^1^. Immunotherapy has revolutionized cancer treatment, with immune checkpoint inhibitors successfully treating several malignancies^2,3^. However, only microsatellite instable (MSI) CRC benefits from immune checkpoint inhibitory approaches, such as targeting CTLA-4 or PD-L1, while clinical responses remain low in microsatellite-stable (MSS) CRC patients^4,5^. This limited efficacy in MSS CRC has been primarily attributed to low mutational burden and lack of immune cell infiltration, resulting in fewer neoantigen targets for either endogenous or αβT cell-based immunotherapies^6–8^. Since the majority of CRC patients are microsatellite stable, identifying new therapeutic targets for this group is of critical importance^9^.

A promising alternative may lie in γδT cells. γδT cells can recognize tumors without the requirement of classical neoantigen presentation via major histocompatibility complex (MHC) molecules and, unlike αβT cells, can respond directly to cellular stress^10^. This makes them attractive candidates for immunotherapy in tumors with a low mutational burden and/or reduced MHC expression. γδT cells are classified by their δ chain usage: Vδ2⁺ γδT cells predominate in circulation, while Vδ1⁺ and Vδ3⁺ γδT cells are enriched in epithelial tissues. In MSS CRC, Vδ1⁺ γδT cells have been reported as either dysfunctional^11^ or tumor-promoting^12^. Yet, in HLA class I-negative MSI CRC, Vδ1⁺ γδT cells play a key role in anti-tumor responses following immune checkpoint inhibition^13^. Further studies have linked the presence of intra-tumoral γδT cell to improved survival, suggesting a generally protective and potentially therapeutic role, regardless of treatment strategy^14,15^. These findings align with studies showing that γδT cell infiltration correlates with a good prognosis in many cancer types^16,17^. In CRC specifically, δ chain usage appears to influence anti-tumor immunity, with Vδ1⁺ and Vδ7⁺ γδT cell populations implicated in early tumor control^18^. Functionally, NKp46⁺ Vδ1⁺ γδT cell subsets have been shown to limit CRC metastasis to the liver in *in vivo* experiments in murine model, reinforcing the importance of co-receptor interactions, such as those mediated by NKp46, in orchestrating effective tumor immunity^19,20^. Despite these promising observations, translation of murine findings to humans remains limited, due to substantial differences between murine and human γδT cell repertoires^21^. Additionally, most transcriptomic studies have lacked robust functional validation of patient-derived γδT cells, largely due to their systemic low abundance, and challenges in *ex vivo* expansion^13^.

Understanding the behavior of γδT cells in solid tumors has been hampered by their rarity across tissue compartments, and by the absence of integrated platforms capable of capturing function, migration, clonality, and transcriptional state in parallel^13–20^. While previous studies have shed light on γδT cell phenotypes and transcriptomic profiles, they have lacked the tools to link tumor reactivity with spatial context, receptor specificity, and migratory behavior. This gap has limited our ability to define the ontogeny and effector potential of tumor-reactive γδT cell subsets across tumor sites. To overcome these limitations, we developed a functional γδT-omics platform that enables the isolation, expansion, and multi-dimensional profiling of γδT cells from primary and metastatic tumor-infiltrating lymphocytes (TILs), lamina propria lymphocytes (LPLs) from the adjacent healthy colon, and peripheral blood lymphocytes (PBLs) across 31 MSS CRC patients. Both bulk-expanded γδT cell populations, and individual clones were analyzed through high-throughput functional screening, biomimetic migration assays, γδTCR repertoire analysis, and transcriptomic profiling. By integrating these data with publicly available single-cell datasets, we identified compartment-specific γδT cell states and uncovered both highly tumor-reactive and functionally impaired subsets, thereby mapping the spatial, clonal, and functional landscape of γδT cells and highlighting key receptors involved in anti-tumor responses (Scheme 1).

**Scheme 1.**
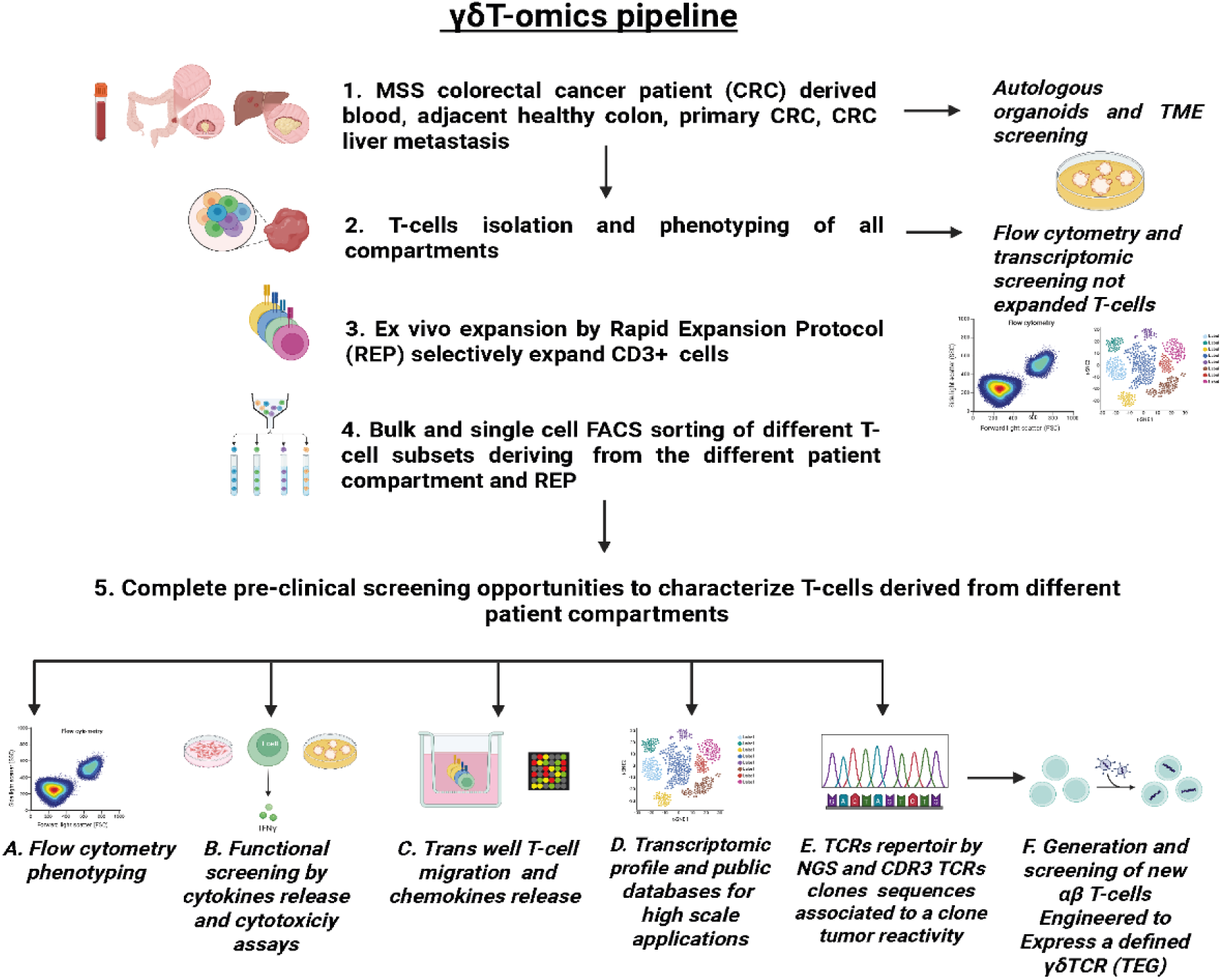
**Experimental γδT-omics pipeline of biopsy isolations in MSS CRC patients and multi-level screening of T-cells deriving from different compartments,** including primary CRC Patient Tumor-infiltrating Lymphocytes (PAT pTILs), CRC liver metastasis Patient tumor-infiltrating Lymphocytes (PAT mTILs), Patient Lamina propria Lymphocytes (PAT LPLs), and both Patient (PAT) and healthy donor (HD) Peripheral Blood Lymphocytes (PBLs).

## Results

### Vδ1^+^ T cells preferentially reside in the primary colorectal tumor site

To characterize the γδT cells immune landscape of in colorectal cancer (CRC) patients, mononuclear cells (MCs) were isolated from primary tumor tissue (MCs containing primary tumor-infiltrating lymphocytes, pTILs), adjacent healthy colon tissue (MCs containing lamina propria lymphocytes, LPLs), and pre-operative peripheral blood samples (peripheral blood mononuclear cells, PBMCs) collected from 31 patients with microsatellite-stable (MSS) CRC undergoing surgical resection (Table 1, Cohort1). Next, patient-derived MCs, deriving from PBMCs, LPLs, and pTILs were analyzed from matched patient samples (PAT), and PBMCs from healthy donors (HD) were used as a reference for comparison (Scheme 1). The MCs density in the peripheral blood of CRC patients (n=28) did not significantly differ from that of healthy donors (n=20) (Figure 1A). However, colon-infiltrating MCs counts differed significantly between unaffected adjacent colon tissue lamina propria mononuclear cells (LPMCs named LPLs, n=20) and primary CRC lesion MCs (named pTILs, n=26), with higher immune cell numbers observed in the tumor tissue. Next, T cell populations from peripheral blood lymphocytes (PBLs), pTILs, and LPLs were phenotypically characterized by flow cytometry. A significantly lower density of γδT cells was observed in patient-derived PBLs compared to healthy donor-derived PBLs, while αβT cell numbers showed no significant differences between groups (Figure 1B–C). Interestingly, both γδ and αβT cell densities were higher in primary CRC tissue, compared to unaffected colon tissue ( Figure 1D–E). The densities of αβ and γδT cells within CRC tumor tissue varied widely between patients. While αβT cell density did not correlate with the maximal tumor diameter at surgical resection, a clear tendency towards negative correlation was observed between γδT cell density and maximal tumor size (r² = 0.45, p = 0.07; Figure 1F-G). The expression of tissue residency markers CD103 and CD69^22^ was elevated in T cells that were isolated from colon tissue, suggesting that colon-infiltrating T cells are resident, rather than circulating (Supplementary Figure 1). Upon examining the densities and proportions of γδT cell subsets across tissue compartments, it was observed that Vδ2⁺ γδT cells dominated in healthy donor PBLs_γδ_, whereas patient-derived PBLs_γδ_ showed a lack of Vδ2⁺ γδT cells compared to the healthy condition (Figure 1H). Instead, the frequency of Vδ1⁺ subset was significantly higher across all patient compartments compared to HD PBMCs (Supplementary Figure 2), with the highest density observed in pTILs_γδ_ compared to other γδT cell subsets and Vδ1⁺ LPLs_γδ_ (Figure 1I).

**Table 1.** Cohorts characterization of patients with CRC. CRC patients cohort 1 used for functional γδT-omics and CRC patient cohort 2 used for clinical validation, were stratified according to sex, age and major clinical manifestation.

| Feature | Variable | CRC patient Cohort 1 (n=31)<br>"Functional $\gamma\delta$T-omics" | CRC patient Cohort 2 (n=69)<br>"Clinical validation" |
| --- | --- | --- | --- |
| Number of MSS patients, n (%) |  | 31 (100) | 69 (100) |
| Gender, n (%) | M | 23 (74.20) | 44 (63.77) |
|  | F | 8 (25.80) | 25 (36.23) |
| Age, mean (±SD) | Mean of the patients (SD) | 65 (±2.36) | 56.30 (±11.30) |
| Primary tumor location, n (%) | Colon | 26 (83.87) | 47 (61.70) |
|  | Rectum | 5 (16.13) | 22 (38.30) |
| Tumor stage, n (%) | 1 | 4 (12.9) | 2 (2.9) |
|  | 2 | 8 (25.8) | 7 (10.14) |
|  | 3 | 11 (35.48) | 36 (52.17) |
|  | 4 | 3 (9.68) | 24 (34.78) |
|  | ND | 5 (16.13) | 0 (0) |
| Metastatic status, n (%) | Non-metastatic | 20 (64.52) | 0 (0) |
|  | Metastatic | 11 (35.48) | 69 (100) |
| Tumor type, n (%) | Adenocarcinoma | 29 (93.55) | 69 (100) |
|  | Medullary Carcinoma | 2 (6.45) | 0 (0) |
| Samples per source of tissue, n (%) | Primary Tumor | 28 (90.32) | 69 (100) |
|  | Liver metastasis | 3 (9.68) | 69 (100) |
|  | Adjacent healthy colon | 21 (67.74) | 0 (0) |
|  | Peripheral Blood | 28 (90.32) | 0 (0) |

**Figure 1.**
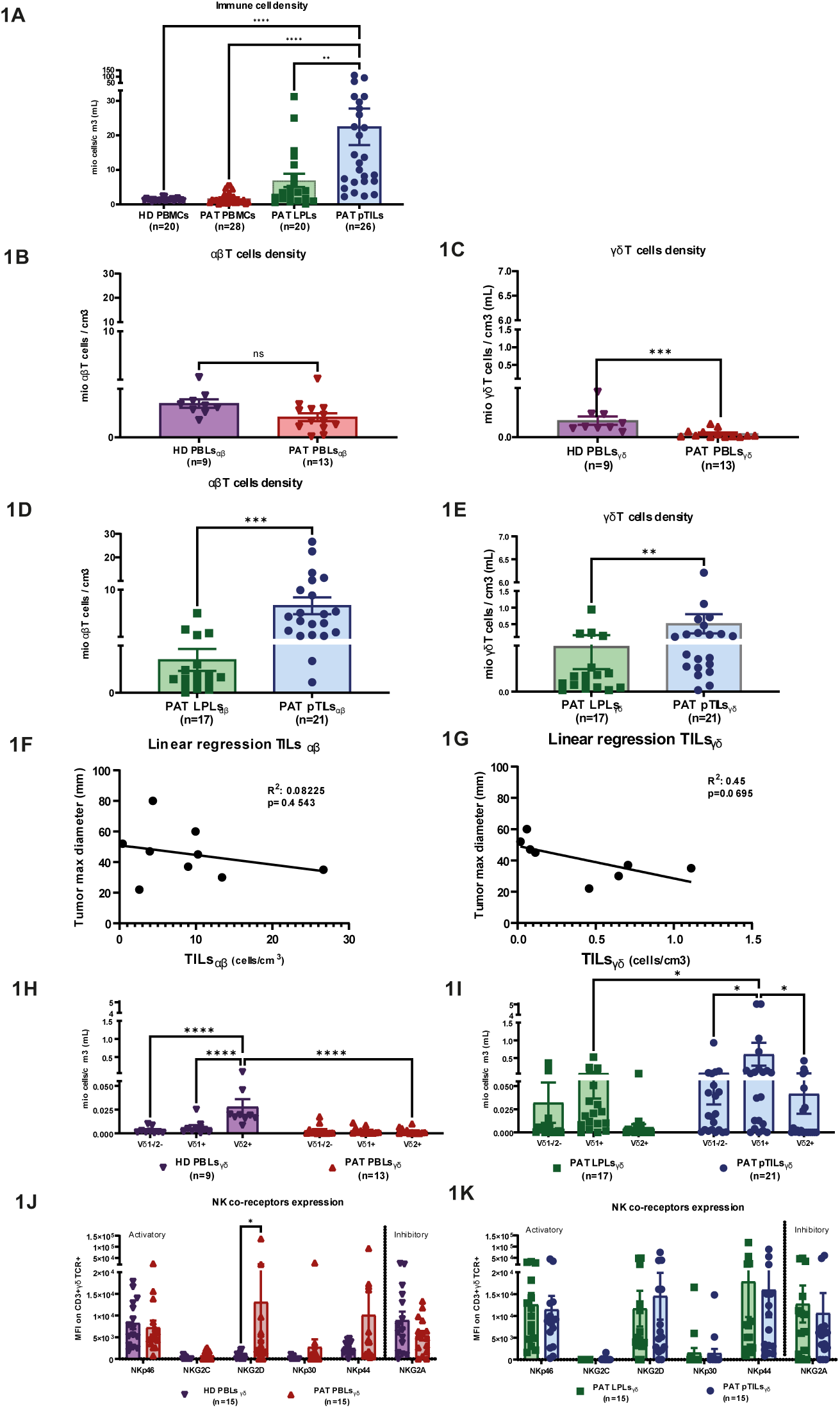
*In vitro* characterization and comparison of γδT cell diversity in CRC patients and HDs. **(A)** Using flow cytometry (Fortessa BD and DIVA software), the diversity and density of different immune cell populations was characterized per cm^3^, across the different biological compartments. Immune cells density was determined in the different biological compartments by normalization of live mononuclear cell counts to the volume of the source material. **(B)** Density of circulating αβT cells and **(C)** Density of γδT cells in HD PBLs and PAT PBLs. **(D)** Levels of αβT cells and **(E)** Density of γδT cells in either healthy colon tissue (PAT LPLs) or primary CRC lesion (PAT pTILs). **(F)** Linear correlation of the maximal tumor diameter and PAT pTILsαβ or **(G)** PAT pTILsγδ. **(H-I)** Concentration of Vδ1-/Vδ2-, Vδ1+, and Vδ2+ T cell populations within the different biological compartments, **(H)** HD and PAT PBLsγδ and colon resident **(I)** PAT LPLsγδ and pTILsγδ. **(J-K)** Utilizing flow cytometry, co-receptor surface expression within the different γδ T cell subsets was determined on both **(J)** Patient (PAT) and healthy donor (HD) Peripheral Blood Mononuclear Cells (PBLsγδ) and **(K)** from Patient Tumor-infiltrating Lymphocytes (PAT pTILsγδ), Patient lamina propria Lymphocytes (PAT LPLsγδ). Co-receptor expression was determined for the total γδT cell population. Not all patients had paired samples. Number of n represents the single patients. Mean values ± SEM are represented with error bars. Statistical significance was either determined by two-way paired t-test **(B-E)** or two-way ANOVA **(A,H,I,J,K)**.

To better understand whether γδT cells exhibit differential characteristics across distinct tissue compartments in CRC patients, various γδT cell subsets were phenotypically analyzed by flow cytometry. The majority of LPLs_γδ_ and pTILs_γδ_ displayed a central memory phenotype (CD45RA⁻ CD27⁺) (Supplementary Figure 3). Considering the important role played by NK co-receptors in γδT cells tumor recognition, surface expression analysis of the most relevant NK receptors were performed. This analysis further revealed that while higher expression of NKG2D was found in patient PBLs_γδ_ compared to HD PBLs_γδ_ (Figure 1J), no differences in NK co-receptors expression was revealed between LPLs_γδ_ and pTILs_γδ_ (Figure 1K). Taken together, these data indicate a dominant presence of Vδ1⁺ γδT cells in all matched tissue samples from MSS CRC patients, with particularly high density within tumor lesions, suggesting a potentially important role in the immune response to CRC.

### *Ex vivo* expanded, Vδ1^+^ pTILs_γδ_ demonstrate the strongest T cell anti-CRC tumor-reactivity, independent of their high NKG2D expression levels

To comprehensively investigate γδT cell reactivity in CRC patients *in vitro,* a rapid expansion protocol (REP)^23^ was employed to expand freshly isolated and FACS-sorted PBLs, pTILs and LPLs derived γδT cell subsets. Vδ1^-^/2^-^, Vδ1⁺, and Vδ2⁺ γδT cell subsets from CRC patients (n=12; labeled “PAT” in figures) and healthy donors (n=7; labeled “HD”) were assessed (Scheme 1). Upon *ex vivo* expansion, surface expression of the activating co-receptor NKG2D was significantly increased in Vδ1^-^/2^-^ pTILs_γδ_ (Figure 2A) and Vδ1⁺ (Figure 2B) γδT cells derived from both pTILs_γδ_ and LPLs_γδ_ were compared to patient-derived PBLs_γδ._ A similar trend (p=0.1348) was observed in NKG2D expression within the Vδ2⁺ subset (Figure 2C) together with high levels of NKG2A expression on patient derived Vδ2⁺ γδT cell, resulting significant for patient Vδ2⁺ PBLs_γδ_ when compared to Vδ2⁺ HD PBLs_γδ_. Next, co-culture of expanded γδT cells with CRC tumor cell lines, Caco-2 and HT-29, revealed that Vδ1^+^ pTILs_γδ_ exhibited the highest numbers of cells releasing IFNγ in response to both tumor lines (Figure 2D; Supplementary Figure 4) and effectively killed HT-29 cells (Supplementary Figure 5). Notably, Vδ1^+^ pTILs_γδ_ also responded to autologous CRC organoids, compared to pTILs_αβ_ (Figure 2E). No correlation between the expression levels of NKG2D and IFNγ production in response to HT-29 was observed (Figure 2F), nor was there a correlation with other tested NK-like receptors (Supplementary Figure 6). Nevertheless, we evaluated the functional impact of NKG2D, by assaying the IFNγ production by tumor-reactive Vδ1^+^ pTILs_γδ_, expressing comparable NKG2D levels (Supplementary Figure 7) in the presence or absence of NKG2D blockade. Reduced tumor reactivity was observed in only 1 out of 5 patients upon NKG2D blocking (Figure 2G), suggesting that NKG2D plays a limited role in mediating tumor reactivity in this γδT cell subset. Vδ1^+^ pTILs_γδ_ exhibited greater cytotoxicity than Vδ1^+^ LPLs_γδ_, a pattern that was not affected by blocking of NKG2D (Figure 2H). In conclusion, *ex vivo* expanded Vδ1⁺ pTILs_γδ_ derived from primary CRC lesions demonstrate enrichment of reactive cells against CRC tumors. This functional capacity appears to be independent of their elevated NKG2D expression, suggesting alternative mechanisms of tumor recognition and cytotoxicity.

**Figure 2.**
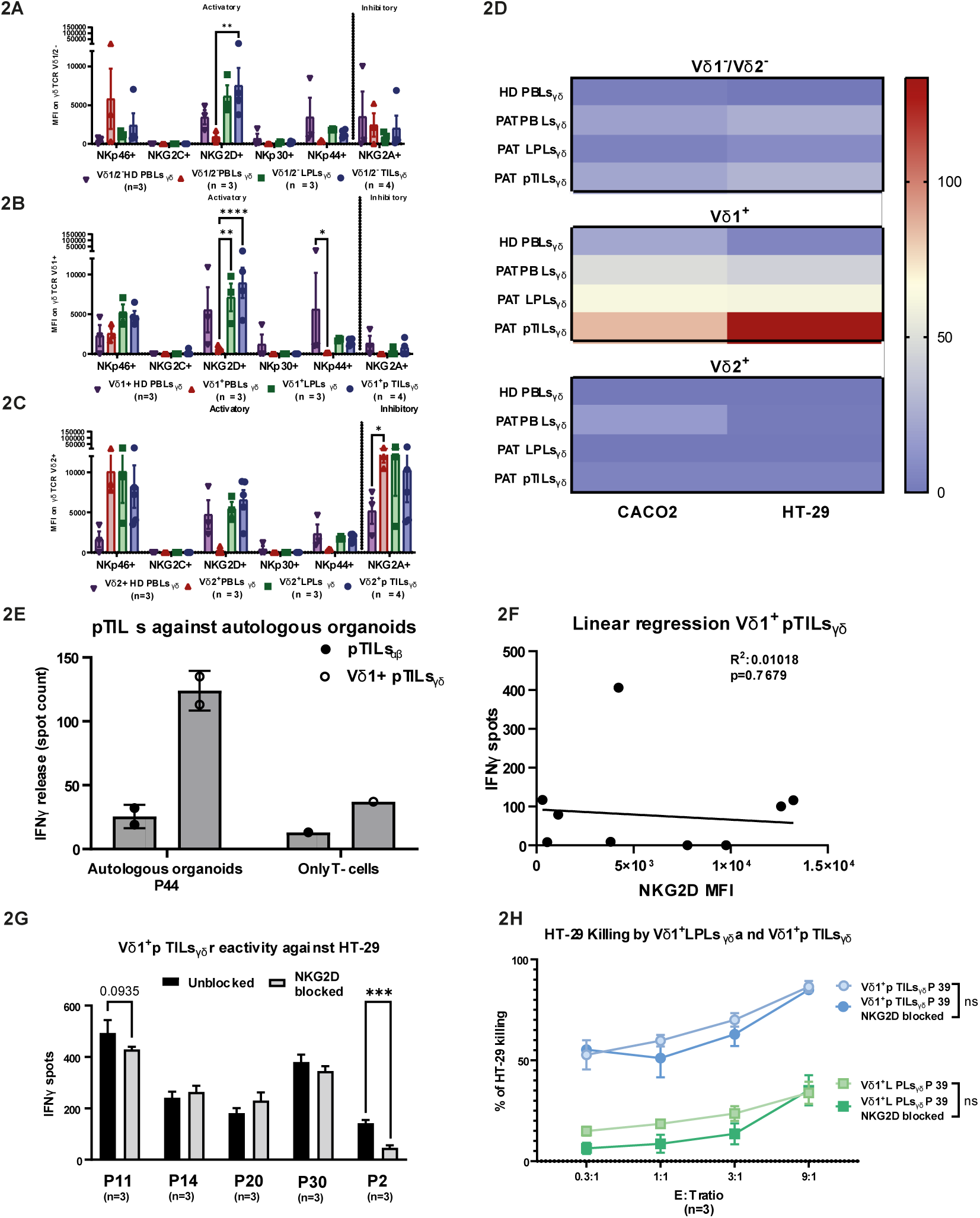
Vδ1+ pTILsγδ exhibit high NKG2D expression and high tumor reactivity, with no observed correlation. Following the pipeline described in scheme 1, γδ and αβ T cell subsets from Patient Tumor-infiltrating Lymphocytes (PAT pTILs), Patient Lamina propria Lymphocytes (PAT LPLs), and both Patient (PAT) and healthy donor (HD) Peripheral Blood Lymphocytes (PBLs). Different T cell populations were expanded *in vitro* through the Rapid Expansion Protocol (REP), and subsequently isolated using FACS bulk sorting. Afterwards, isolated T cell populations from the different comparments were expanded for various cycles of REP to produce sufficient cell numbers for co-receptor expression analysis and functional testing. Co-receptor expression patterns were determined by flow cytometry for **(A)** the Vδ1^-^/2^-^ HD PBLsγδ and PAT PBLsγδ/LPLsγδ/pTILsγδ subset, **(B)** Vδ1^+^ HD PBLsγδ and PAT PBLsγδ/LPLsγδ/pTILsγδ subset and **(C)** the Vδ2^+^ HD PBLsγδ and PAT PBLsγδ/LPLsγδ/pTILsγδ subset. **(D)** IFNγ release by either Vδ1^-^/Vδ2^-^, Vδ1^+^, or Vδ2^+^, from the different sources, HD PBLsγδ and PAT PBLsγδ/LPLsγδ/pTILsγδ, in response to co-culture with HT-29 and Caco-2 cell lines. Tumor reactivity was assessed after a 24-hour co-culture of T cells with CRC cell lines at a 1:3 ratio, normalized to T cell only conditions, using an IFNγ ELISpot assay. **(E)** IFNγ release by either pTILsαβ or Vδ1+ pTILsγδ bulks in response to co-culture with autologous organoids. Tumor reactivity was assessed after a 24-hour co-culture of T cells with CRC organoids at a 1:3 ratio, T cell only condition has been shown as a negative control, using an IFNγ ELISpot assay. **(F)** Linear regression between the NKG2D expression of the expanded Vδ1+ pTILsγδ and IFNγ release against HT-29. **(G)** IFNγ release of the expanded Vδ1+ pTILsγδ as performed in **(E)** but with or without a one hour pre-incubation with an NKG2D blocking antibody (10ug/mL). **(H)** Luciferase based killing by Vδ1+ LPLsγδ and pTILsγδ with or without a one hour pre-incubation with either an NKG2D blocking antibody (10ug/mL) targeting the CRC cell line HT-29 at multiple E:T ratios, normalized to the target only condition. Mean values ± SEM are represented with error bars. Statistical significance was determined by either a one-way ANOVA **(A-C)**, two-way ANOVA **(D)**, or one-way t-test **(G,H).**

### Vδ1^+^ pTILs_γδ_ are more frequent in tumors and exhibit higher potency against CRC than other TIL subsets

To substantiate our findings at the clonal level, and further investigate the molecular mechanisms of tumor recognition by tumor-infiltrating Vδ1⁻/2⁻ and Vδ1⁺γδT cells, nearly 9,000 single Vδ2⁻ γδT cell clones were isolated by FACS from PBLs of healthy donors and CRC patients, as well as from LPLs and pTILs of 11 CRC patients following Scheme 1. On average, approximately 10% of these single cells expanded to ≥10⁴ cells using a rapid expansion protocol (REP) (Supplementary Figure 8), and 133 clones reached 0.2 × 10⁶ cells, enabling further phenotypic and functional characterization. The expanded clones were functionally tested by quantifying IFNγ release following co-culture with the CRC cell lines HT-29 and Caco-2. Subsequently, the frequency of tumor-reactive Vδ1^-^/2^-^ γδT cell clones was calculated for each CRC patient, and healthy donor compartment. No statistically significant differences in tumor-reactive Vδ1^-^δ2^-^ clone frequencies were observed between patient-derived tissue compartments and healthy donors (Figure 3A). In contrast, the frequency of tumor-reactive Vδ1⁺ γδT cell clones derived from pTILs was significantly higher (67%) than those from other compartments, including patient-derived Vδ1^+^ LPLsγδ (33%) and PBLs_γδ_ (22%), and healthy donor-derived PBLs_γδ_ (16%) (Figure 3B).

**Figure 3.**
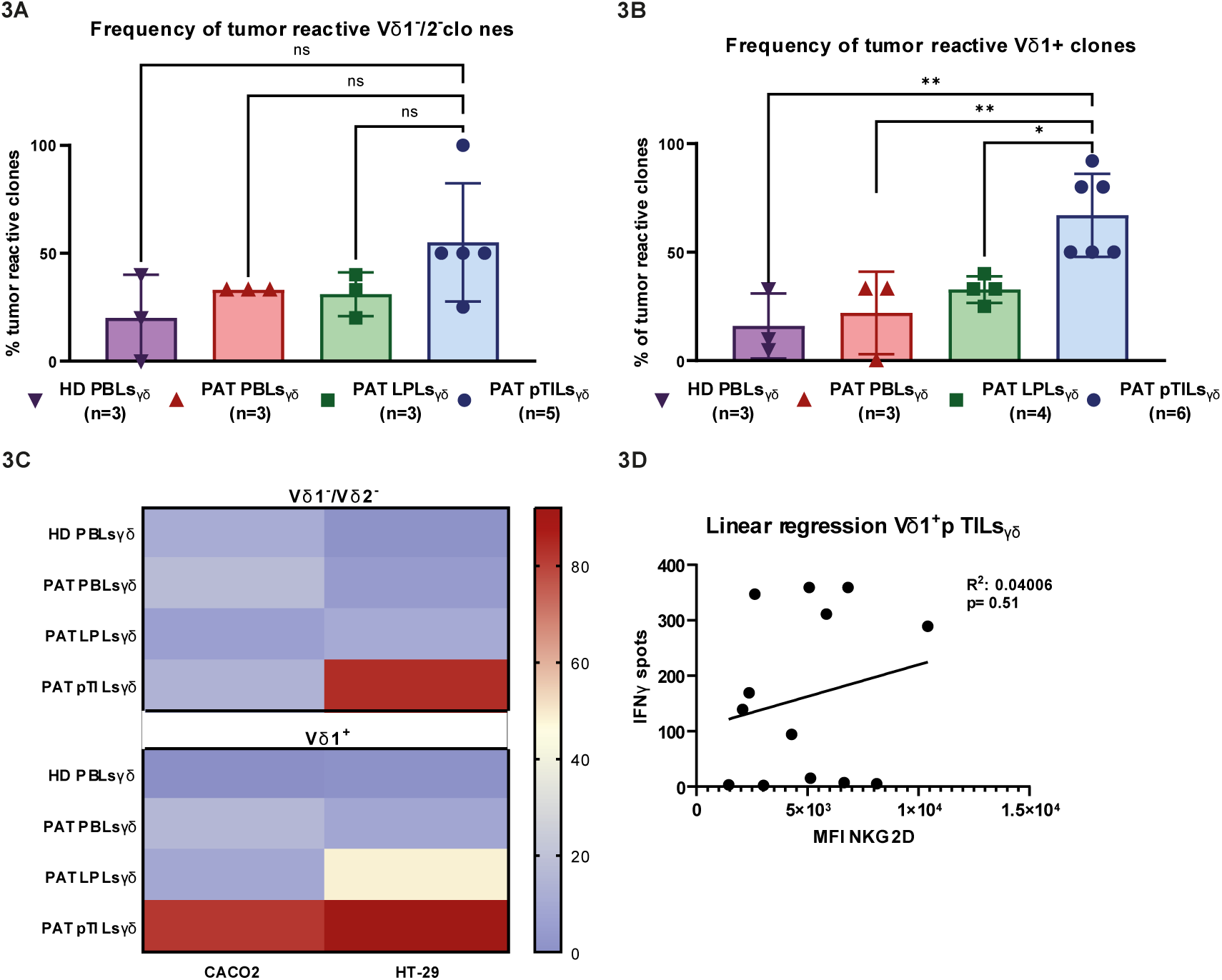
High tumor reactivity and increased frequency of tumor-reactive Vδ1+ pTILsγδ clones. Following the experimental pipeline described in scheme 1, single γδ T cell subsets from Patient Tumor-infiltrating Lymphocytes (PAT pTILsγδ), Patient Lamina propria Lymphocytes (PAT LPLsγδ), and both Patient (PAT) and healthy donor (HD) Peripheral Blood Lymphocytes (PBLsγδ) were single cell sorted, and expanded for various cycles of REP to produce sufficient cell numbers for co-receptor expression analysis and functional testing. **(A)** Frequency of tumor-reactive Vδ1-/ Vδ2 or **(B)** Vδ1+ T cell clones derived from the different biological compartment, PBLsγδ/LPLsγδ/pTILsγδ. Tumor reactivity was determined by the production of 30 IFNγ spots after a 24-hour co-culture of T cells with one of the CRC cell lines at a 1:3 ratio, normalized to T cell only conditions. **(C)** IFNγ release by either Vδ1+ or Vδ1-/Vδ2-expanded γδT cell bulks derived from the different sources, in response to co-culture with HT-29 and Caco-2 cell lines. Tumor reactivity was assessed after a 24-hour co-culture of T cells with CRC cell lines at a 1:3 ratio, normalized to T cell only conditions, using an IFNγ ELISpot assay. Each dot represents an individual clone. **(D)** Linear regression between the NKG2D expression of the expanded Vδ1+ pTILsγδ clones and IFN-γ release against HT-29. Individual data points indicate values from individual CRC patients of HDs (**A,B)** or individual T cell clones **(C,D)**. Not all patients had paired samples. Mean values ± SEM are represented with error bars. Statistical significance was determined by ordinary one-way ANOVA for the different subsets.

Assessment of individual clone activation and consequent IFNγ release revealed that 5 out of 9 Vδ1^-^/2^-^ pTILs_γδ_ clones were sensitive to activation in response to HT-29, with spot counts ranging from 84 to 244 (Figure 3C, Supplementary Figure 9). However, this was lower than the response observed in Vδ1^+^ pTILs_γδ_ clones, of which 16 out of 25 clones were producing IFNγ, with spot counts ranging from 55 to 359 (Figure 3C, Supplementary Figure 9). These results confirm that tumor reactive Vδ1⁺ γδT cells are enriched within CRC lesions. Phenotypic analysis of tumor-reactive Vδ1^+^ pTILs_γδ_ clones showed that consistent with polyclonal observations (Figure 2F), high IFNγ releasing Vδ1^+^ pTILs_γδ_ were not associated with increased expression of the activating co-receptor NKG2D on their surface (Figure 3D). In conclusion, while tumor-reactive Vδ1⁻/2⁻ and Vδ1⁺ γδT cell clones were present in healthy donor-derived PBLs, as well as in patient-derived PBLs and LPLs, the highest frequency of tumor reactivity per patient was found among the Vδ1⁺ γδT cell clones from pTIL, as evidenced by elevated IFNγ producing cells in response to CRC tumor targets.

### Defined Vδ1^+^ γδTCR transfer into engineered αβT cells preserves tumor-reactivity

To characterize the role of the γδTCR in tumor recognition, the γ and δTCR gene sequences from two tumor-reactive (clones 09TP1G1 and 04TP4C4), and two non-tumor-reactive (clones 04TP2F4 and 14TP2C3) Vδ1⁺ single-cell clones derived from pTILs_γδ_ were isolated and used to generate αβT cells receptor-engineered to express a defined γδTCR instead of their native αβTCRs (TEGs)^24^ in CD4^+^ αβT cells (Figure 4A). Since CD4⁺ αβ T cells naturally express low levels of NK-like surface co-receptors, this model allows exclusive evaluation of γδTCR tumor recognition^25^. Co-culturing the original T cell clones or their corresponding TEGs with a panel of tumor cell lines, including CRC cell line HT-29, revealed that γδTCRs from tumor-reactive clones 04TP4C4 and 09TP1G1 mediated robust counts of IFNγ releasing cells, comparable to, or exceeding those of the parental γδT cell clones (Figure 4B,C). In contrast, γδTCRs from non-tumor-reactive clones (04TP2F4 and 14TP2C3) induced minimal IFNγ production (Figure 4D, E). These findings demonstrate that γδTCR is a primary driver of tumor recognition, potentially modulated by co-receptor activity in the parental γδT cells. Furthermore, IFNγ release by tumor reactive 04TP4C4, 09TP1G1 TEGs, but not Vδ2^+^ γδTCR TEG-LM1^24^, was observed across a broader panel of tumor cell lines, including glioma (U87), head and neck carcinoma (SCC9), multiple myeloma (RPMI-8226), and acute myeloid leukemia (ML-1), in addition to HT-29 (Supplementary Figure 10). This suggests that γδTCR-mediated reactivity is not limited to CRC. In line with this, direct tumor-killing activity was confirmed against HT-29, RPMI-8226, and SCC9 (Figure 4F; Supplementary Figures 11A and 11B). Taken together, the tumor-reactivity of Vδ1^+^ pTILs_γδ_ clones was preserved, following their γδTCR transfer into αβT cells with low expression of additional innate co-receptors. This finding implies that the γδTCR derived from Vδ1^+^ pTILs_γδ_ drives tumor recognition and response to tumor antigens in CRC.

**Figure 4.**
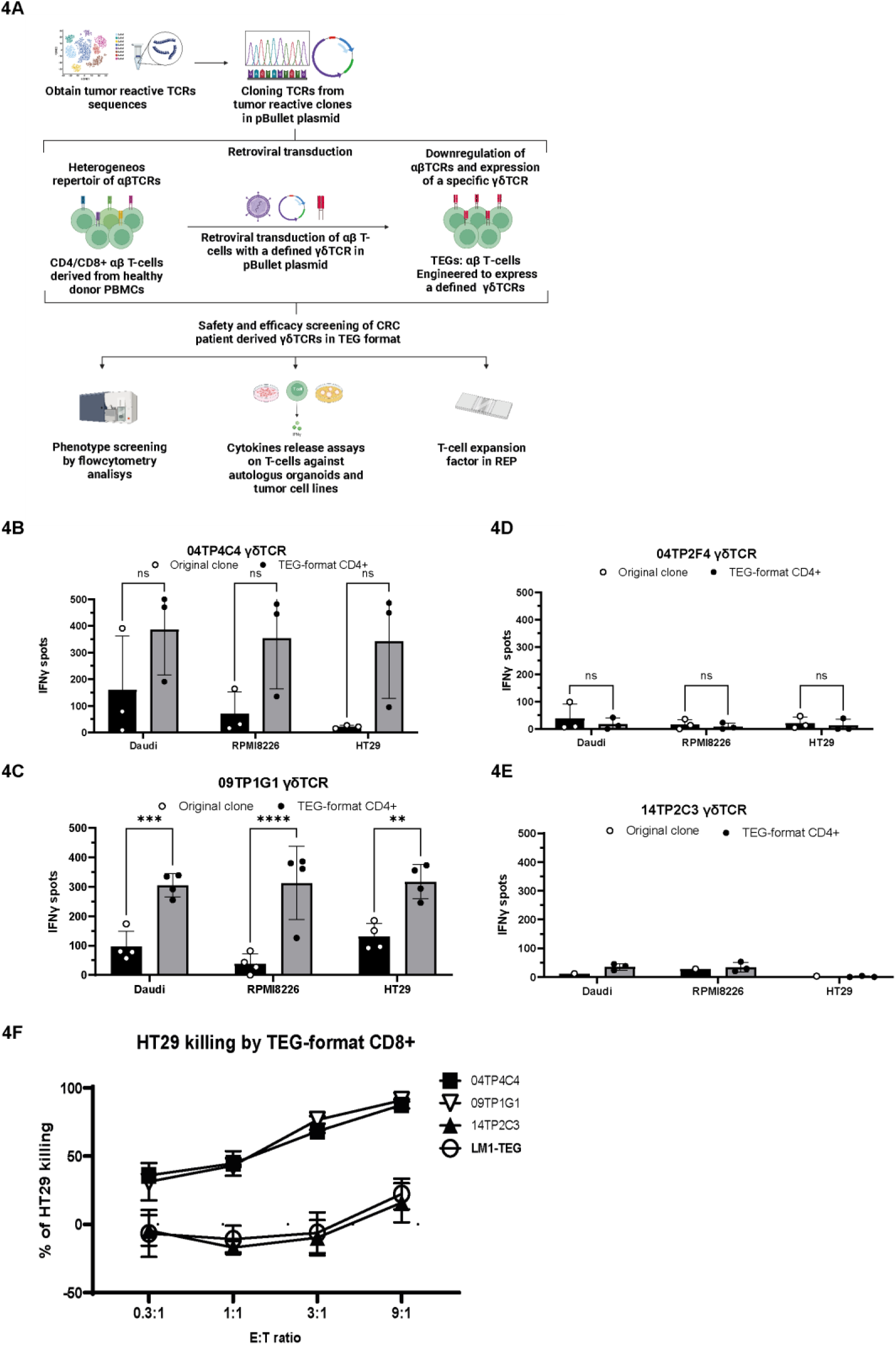
High and broad tumor reactivity upon transfer of Vδ1+ TCRs from tumor reactive clones to αβ T cells. **(A)** Experimental pipeline: αβ T cells were isolated from HD PBMCs and transduced with the γδ TCRs from the PAT pTILs derived Vδ1+ T cell clones according to the previously published protocol. The transduced T cells (TEGs) were expanded according to the REP and assessed for cytotoxic activity and cytokine production in functional assays, next to their original clone. Two TCRs were isolated from the tumor reactive clones **(B)** 04TP4C4, **(C)** 09TP1G1 and two TCRs were derived from the non-tumor reactive clones **(D)** 04TP2F4, **(E)** 14TP2C3. IFN-γ release by either the Vδ1+ clone or corresponding CD4+ TEGs in response to co-culture with the tumor cell lines Daudi, RPMI-8226 and HT-29. Tumor reactivity was assessed after a 24-hour co-culture of T cells with the tumor cell lines at a 1:3 ratio, using an IFN-γ ELISpot assay, and normalized to a no target control. **(F)** Luciferase based killing by the Vδ1+ V09TP1G1 and 04TP4C4 TEGs and the Vδ2+ TEG001 and mock TEG-LM1cell targeting the CRC cell line HT-29 at multiple E:T ratios, normalized to the target only condition. Mean values ± SD are represented with error bars. Statistical significance was determined (B-E) by two-way ANOVA.

### Vδ1^+^ LPLs_γδ_ have the highest T cell migration capacity to CRC tumor cells

To assess the migratory potential of Vδ1⁺ γδT cells from various compartments toward the tumor site, expanded bulk Vδ1⁺ γδT cells derived from patient PBLs, LPLs, or pTILs were subjected to a 3D trans-well migration assay in the presence of the CRC cell line Caco-2 (Figure 5A). Among the tested subsets, Vδ1^+^ LPLs_γδ_ exhibited significantly greater migration toward the tumor cells, compared to either Vδ1^+^ TILs_γδ_ or Vδ1^+^ PBLs_γδ_ (Figure 5B) suggesting that LPLs_γδ_ possess the highest intrinsic migratory capacity toward CRC lesions in patients. In contrast, the reduced migration observed in Vδ1^+^ pTILs_γδ_ may indicate a loss of migratory capacity upon residency within the TME. To examine whether there is a role for the γδTCR specificity in the observed tumor-directed migration, TEGs were tested in the same trans-well system. Both tumor-reactive Vδ1^+^ 04TP4C4 TEGs and the existing literature on Vδ2^+^ A3-TEG^26^, showed enhanced migration toward the tumor site compared to non-reactive Vδ1^+^ 14TP2C3 TEGs, or the mock Vδ2^+^ TEG-LM1^24^ (Supplementary Figure 12). In conclusion, these findings suggest that Vδ1^+^ LPLs_γδ_ possess strong migratory potential, and are likely contributors to the pool of tumor-reactive Vδ1⁺ clones observed in pTILs (Figure 3B). However, once these LPLS-derived γδT cells enter the TME, their migration capacity appears to diminish. This highlights that while γδTCR-mediated tumor recognition facilitates the accumulation of γδT cells within the tumor, probably by the upregulating chemoattractants secretion^27^, it is not the sole determinant. The TME itself likely imposes regulatory effects that shape the behavior of infiltrating immune cells. Further investigation is required to better understand the specific actions by both tumor and T cells to modulate tumor infiltration and recognition.

**Figure 5.**
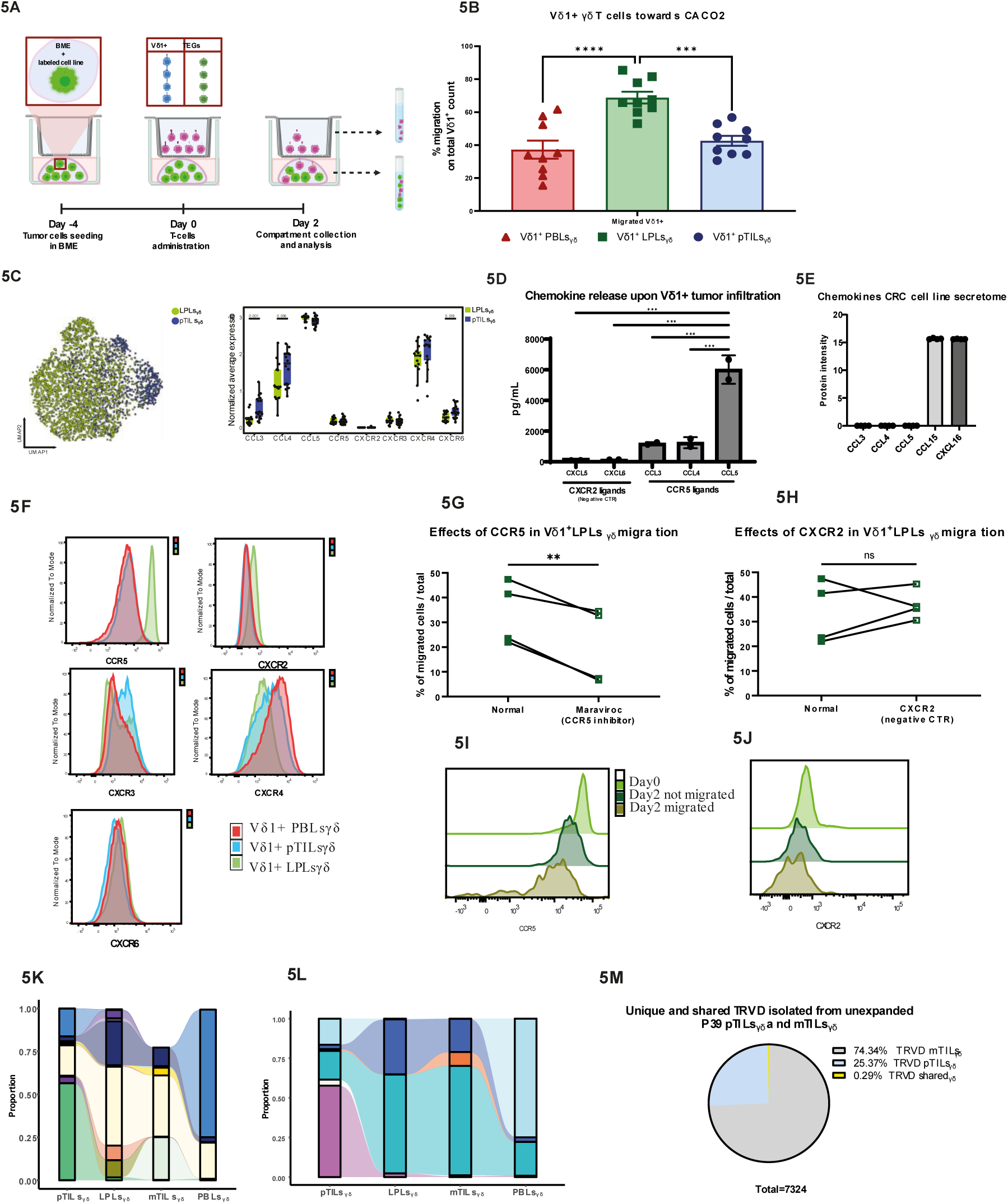
Lamina propria Lymphocytes are the primary source of tumor-reactive. Vδ1+ pTILsγδ **and their migration is dependent on the CCR5 expression.** Schematic timeline of the trans well, ECM experiments. CACO-2 tumor cells were dyed and cultured for four days in Basement membrane extract (BME). Subsequently, different T cell subsets (CD3+ in B, Vδ1+ in C and TEGs in D) were administered in the upper compartment of the trans well, and co-cultured for two days before isolation and analysis of both the non-migrated (upper compartment) and migrated (lower compartment) T cell population. **(B)** Percentage of migrated Vδ1+ γδT cells of total Vδ1+ γδT cells isolated at day 2 post T cell administration of Vδ1+ γδT cell bulks deriving from different biological sources, pTILsγδ, LPLsγδ, PBLsγδ, of the same CRC patient. **(C)** left panel: UMAP representation of γδT cells isolated from the primary CRC (pCRC) dataset comprised of primary colon tumor tissues (pTILsγδ, n=16) and adjacent normal colon tissues (pLPLsγδ, n=13) of MSS CRC patients. Each dot represents a single cell. Right panel: Average normalized gene expression (per compartment) of several key chemokine and chemokine receptors. **(D)** Chemokine release in the supernatant of the trans well lower compartment upon tumor infiltration of LPLsγδ and pTILsγδ **(E)** Chemokine release by CRC cell line upon 48h in cell-conditioned media. n=4, every value represents a single well. **(F)** The surface expression of CCR5, CXCR2, CXCR3, CXCR4 and CXCR6 were tested by flow cytometry in the expanded patient derived Vδ1^+^ LPLsγδ showing strong differences between patient compartments, in particular for CCR5 **(G-H)** Percentage of migrated Vδ1^+^ LPLsγδ of total Vδ1+ T cells isolated at day 2 post T cell administration of Vδ1^+^ bulks either in presence or absence of the CCR5 inhibitor Maraviroc **(G)** and CXCR2 neutralizing antibody **(H)**. **(I-J)** Histogram representing the MFI of CCR5 **(I)** and CXCR2 **(J)** on Vδ1+ LPLsγδ at different time points of the migration assay: day0, day2 not migrated fraction and day2 migrated fraction. Showing a decrease of CCR5 MFI on Vδ1^+^ LPLsγδ upon migration.**(K)** The TRVD1 clonotypes of expanded patient derived Vδ1 bulks originating from pTILs across other compartments generated by Next Generation Sequencing. **(L)** TRVD clonotypes (amino acid sequences) that are shared across all compartments (Vδ1^+^ pTILsγδ, LPLsγδ, mTILs γδ, PBLs γδ). (**M)** 10X 5’ scRNA sequencing analysis on the TRVD repertoire of unexpanded primary and metastatic TILsγδ of the same patient 39 show only the 0.29% of overlap between the two TRVD repertoires. Mean values ± SD are represented with error bars. Statistical significance was determined by (C-D) Ordinary one-way Analysis of Variance (ANOVA), (F-G) Paired T-test.

### Migration of Vδ1^+^ LPLs_γδ_ toward the tumor and their retention at the tumor site is initiated by CXCL16 and regulated by the chemokine receptor CCR5

Since chemokines and their receptors play a pivotal role in T cell migration, we aimed to investigate whether the differential migratory capacity observed between Vδ1⁺ LPLs_γδ_ and pTILs_γδ_ could be attributed to distinct chemokine, or chemokine receptor expression profiles. To this end, a publicly available single-cell RNA sequencing (scRNA-seq) dataset of MSS CRC samples (pCRC dataset), comprising primary colon tumor tissues (n=16) and adjacent normal tissues (n=13), was analyzed^28^. γδT cells were computationally extracted based on original cell annotations, and their expression patterns of key chemokines and chemokine receptors involved in cell migration were assessed (Figure 5C). Transcriptomic analysis revealed similar expression levels of CXCR3 and CXCR4 between compartments, while CXCR2 was not expressed. Notably, CXCR6 expression was significantly higher in pTILs_γδ_ compared to LPLs_γδ_ (Figure 5C). Among the chemokine ligands, high expression of the CCR5 ligands, CCL3, CCL4, and CCL5, was observed in both compartments, with CCL3 and CCL4 significantly elevated in pTILs_γδ_ CCL5 was highly expressed in both pTILs and LPLs populations (Figure 5C). These transcriptomic findings were supported by high protein-level measurements of CCR5 ligands, particularly CCL5, in the supernatants from the lower chamber of the trans-well assay, following tumor infiltration by pTILs_γδ_ and LPLs_γδ_ (Figure 5D). In contrast, CXCL5 and CXCL6, ligands for the non-expressed CXCR2, were undetectable (Figure 5D). Therefore, an unbiased proteomic-based secretome analysis of a representative CRC-tumor cell line was performed. Among the identified 2583 proteins, the only two chemokines found were the ligand for CCR1/3, CCL15, and the ligand for CXCR6, CXCL16,while none of the CCR5 ligands, CCL3, CCL4, CCL5, were detected (Figure 5E). These data suggest that γδT cells are initially attracted to tumor cells via the CXCL16-CXCR6 axis. Upon tumor contact, activation via the γδTCR induces CCL5 secretion to further recruit Vδ1⁺ pTILs_γδ_ to the tumor site. Although CCR5 gene expression was relatively low and comparable between compartments, surface expression profiling in expanded Vδ1⁺ γδT cells revealed markedly higher CCR5 protein levels on LPLsγδ compared to patient-derived pTILs_γδ_ and PBL_γδ_, whereas other tested chemokine receptors were not strongly expressed in LPLs_γδ_ (Figure 5F). To directly test the role of CCR5 in γδT cell migration, trans-well assays were conducted in the presence or absence of the clinically approved CCR5 inhibitor Maraviroc, as well as a neutralizing antibody against the non-expressed CXCR2 (negative control). Blocking of CCR5 significantly reduced the migration of Vδ1⁺ LPLs_γδ_ (Figure 5G), while the CXCR2 blockade had no effect (Figure 5H), confirming CCR5 as a key mediator of Vδ1⁺ LPLs_γδ_ migration towards tumor cells. Additionally, CCR5 surface expression was monitored during migration assays and found to be reduced in migrated Vδ1⁺ LPLs_γδ_ compared to their non-migrated counterparts (Figure 5I), consistent with receptor internalization upon ligand engagement. No changes were observed in CXCR2 (Figure 5J). Hence, our data clarifies whether the detected CCL5, or are produced by Vδ1⁺ γδT cells that migrated towards the tumor and were subsequently activated in the lower chamber. Our findings identify CCR5 as a key regulator of the enhanced migratory capacity and retention of Vδ1⁺ LPLs_γδ_ within CRC tumors, thereby promoting the accumulation of tumor-reactive γδT cells in the TME.

### Vδ1^+^ LPLs_γδ_ are the main source of Vδ1+ pTILs_γδ_ leading to a unique TRVD1 repertoire compared to liver mTILs

To test that Vδ1+ pTILs_γδ_ primarily originate from Vδ1^+^ LPLs_γδ_, a TRVD1 repertoire analysis was performed on patient-derived Vδ1^+^ PBLs_γδ_, LPLs_γδ_, pTILs_γδ_ and additionally Vδ1^+^ T cells isolated from CRC liver metastasis of the same patient (mTILs). In line with the high tumor-directed migratory capacity of Vδ1^+^ LPLs_γδ_ (Figure 5B), most TRVD1 clonotypes, including some unique TRVD1 colon-resident clonotypes, were shared between Vδ1^+^ LPLs_γδ_ and Vδ1^+^ pTILs_γδ_ (Figure 5K), providing strong evidence that Vδ1^+^ LPLs_γδ_ serve as the primary source of the highly tumor-reactive Vδ1^+^ pTILs_γδ_. In contrast, the TRDV1 repertoire of Vδ1^+^ mTILs_γδ_ showed distinct clonotypes that overlapped primarily with patient-derived Vδ1^+^ PBLs_γδ_ and, to a lesser extent, with Vδ1^+^ LPLs_γδ_, but had limited overlap with the Vδ1^+^ pTILs_γδ_ repertoire (Figure 5K). These findings suggest that Vδ1⁺ γδT cells infiltrating metastatic lesions originate from a different compartment, likely the circulation or adjacent liver^29^, rather than from colon-resident Vδ1^+^ LPLs_γδ_. Across all tissues, only six TRDV1 clonotypes were shared, with varying levels of clonal expansion (Figure 5L). The strongest overlap was again observed between Vδ1^+^ LPLs_γδ_ and pTILs_γδ_, while overlap between Vδ1^+^ pTILs_γδ_ and mTILs_γδ_ repertoires was minimal (Figure 5L). In addition, single-cell RNA/γδTCR analysis of Vδ1^+^ pTILs_γδ_ and mTILs_γδ_ from the same patient, before *ex vivo* expansion, confirmed the limited overlap of TRVD clonotypes between the two tumor sites, with only 0.29% of total TRVD sequences shared between pTILs_γδ_ and mTILs_γδ_ (Figure 5M). These results, combined with the previous migration data, point to Vδ1^+^ LPLs_γδ_ as the main source of Vδ1^+^ pTILs_γδ_. The Vδ1^+^ mTILs_γδ_ repertoire varied markedly from that of Vδ1^+^ pTILs_γδ_, suggesting tissue-specific Vδ1^+^ γδT cell repertoires that do not substantially migrate across compartments during metastatic progression.

### Vδ1^+^ mTILs_γδ_ lack tumor-reactivity

Since the Vδ1⁺ TILs_γδ_ TCR repertoires differed between primary tumors and liver metastasis, an investigation into whether these distinct repertoires exhibit different intrinsic capacities to recognize CRC was conducted. To this end, Vδ1^+^ mTILs_γδ_ were isolated from liver metastases of three CRC patients, expanded, and functionally characterized. Subsequently, Vδ1⁺, Vδ2⁺, and Vδ1^-^/2^-^ γδT cell subsets from unmatched pTILs_γδ_ and mTILs_γδ_ were tested for tumor reactivity using IFNγ ELISpot assays, following co-culture with the CRC cell line HT-29. Strikingly, all γδT cell subsets derived from mTILs_γδ_ displayed reduced tumor-reactivity compared to their counterparts from pTILs_γδ_ and significantly lower compared to Vδ1⁺ pTILs_γδ_ (Figure 6A). To further validate this observation, a comparison between the tumor recognition capacity of Vδ1^+^ mTILs_γδ_ and mTILs_αβ_ from patient 31, against autologous CRC organoids was performed. Vδ1^+^ mTILs_γδ_ failed to recognize the autologous tumor target, confirming their loss of tumor reactivity (Figure 6B). Furthermore, the aim was to assess whether this loss of reactivity was due to altered ligand expression on metastatic versus primary tumor cells, or to intrinsic changes in the Vδ1^+^ mTILs_γδ_. IFNγ ELISpot assays using Vδ1^+^ pTILs_γδ_ co-cultured with either autologous primary or metastatic CRC organoids revealed no difference in tumor recognition (Figure 6C), indicating that the tumor express ligands for Vδ1⁺ γδT cell. In conclusion, the observation of diminished tumor recognition observed in Vδ1^+^ mTILs_γδ_ could be associated with the absence of tumor-reactive γδTCRs, consistent with the different repertoire data compared to Vδ1^+^ pTILs_γδ_, and possibly with additional phenotypic alterations acquired within the metastatic microenvironment.

**Figure 6.**
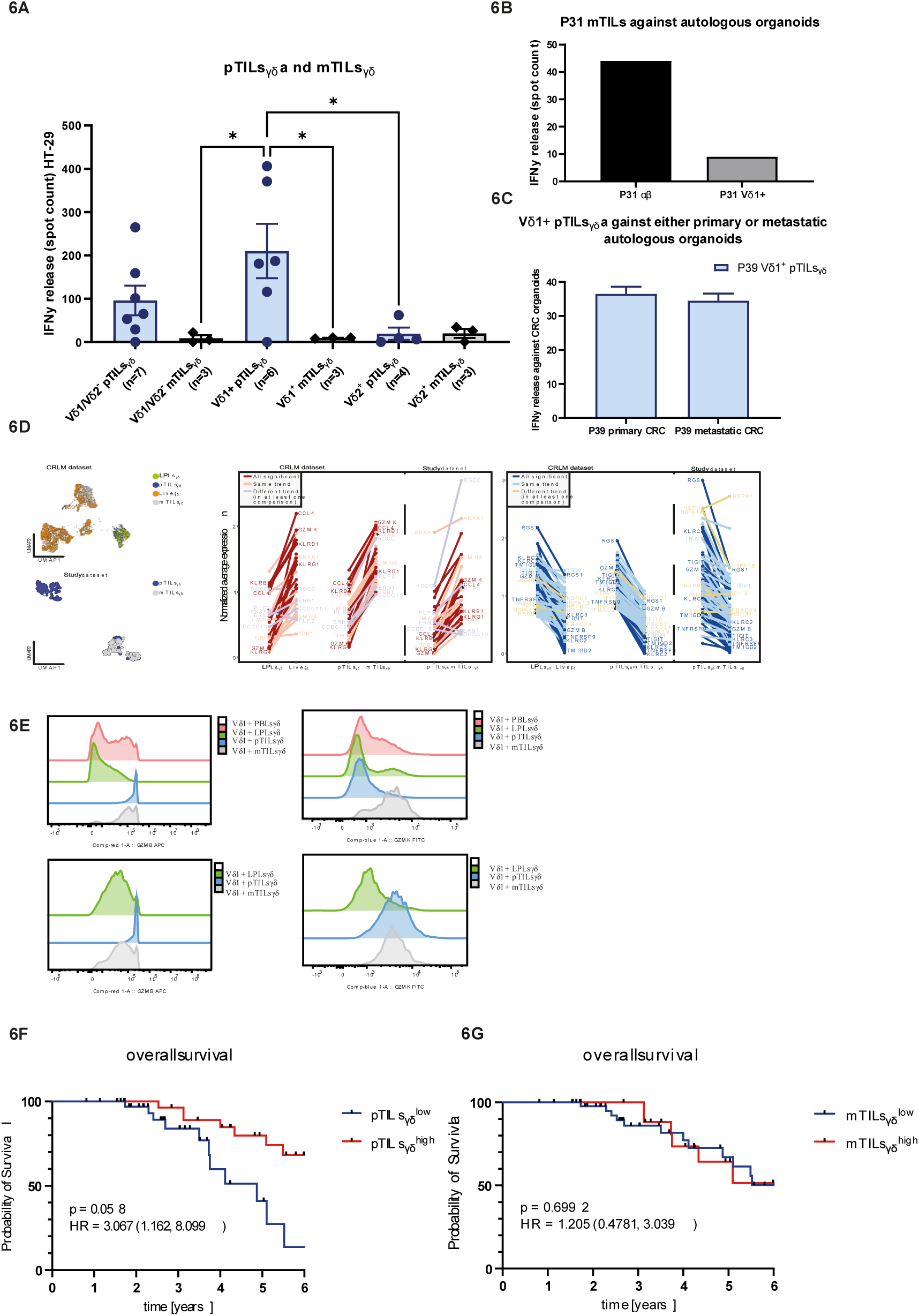
The colon-resident tumor-reactive Vδ1+ T cells do not reach the metastasized lesions and show differences in transcriptomic profile and functionality. **(A)** IFNγ release either Vδ1+, Vδ2+, or Vδ1-/Vδ2-either pTILsγδ or mTILsγδ upon a co-culture with the CRC cell line HT-29. Tumor reactivity was assessed after a 24-hour co-culture of T cells and tumor cells at a 1:3 ratio, using an IFNγ ELISpot assay. Every dots represent a biological replicate. The different TRVD1 repertoires found in pTILsγδ and mTILsγδ resulted in a lower IFNγ release of mTILsγδ bulks against CRC cell lines when compared to pTILsγδ. **(B)** IFNγ release by either mTILsαβ or Vδ1+ mTILsγδ bulks in response to co-culture with autologous organoids. Tumor reactivity was assessed after a 24-hour co-culture of T cells with CRC organoids at a 1:3 ratio, using an IFNγ ELISpot assay. **(C)** ) IFNγ release by Vδ1+ pTILsγδ bulks in response to co-culture, with either primary or metastatic autologous organoids. Tumor reactivity was assessed after a 24-hour co-culture of T cells with CRC organoids at a 1:3 ratio using IFNγ ELISpot assay. **(D)** Top left panel: UMAP representation of γδT cells from the primary CRC and liver metastatis public dataset (GSE164522), named CRLM dataset, colored by tissue of origin (LPLsγδ, n=8; pTILsγδ, n=7; Liverγδ, n=9; mTILsγδ, n=7). Bottom left panel: UMAP representation of scRNA-seq profiles of γδT cells isolated from a single patient (PAT39), derived from primary colon tumor tissue (pTILγδ) and liver metastasis tissue (mTILγδ). Middle and right panels: Averaged normalized gene expression of differentially expressed genes in pTILsγδ vs mTILγδ comparison of the public dataset (GSE164522). **(E)** FACS analysis of the differentiated expressed GZMB and GZMK in expanded Vδ1+ bulks allowed us to confirm a different transcriptomic profilebetween compartments, which remains stable upon ex vivo expansion. **(F)** Kaplan-Meier survival curve of patients with UICC stage IV MSS CRC (n=69) stratified according to the extent of pTILsγδ infiltration into the infiltrative margin of primary tumors. The extent of pTILsγδ infiltration is classified as low for pTILsγδ infiltration <= 10 pTILsγδ per mm2, and classified as high for pTILsγδ infiltration > 10 pTILsγδ per mm^2^. **(G)** Kaplan-Meier survival curve of patients with UICC stage IV MSS CRC (n=69) stratified according to the extent of mTILsγδ infiltration into the infiltrative margin of liver metastatic tumors. The extent of mTILsγδ infiltration is classified as low for mTILsγδ infiltration <= 5 mTILsγδ per mm^2^ and classified as high for mTILsγδ infiltration > 5 mTILsγδ per mm^2^. Genes are color-coded to indicate whether they are significantly differentially expressed, show a consistent expression trend across all comparisons, or exhibit divergent trends (e.g., upregulated in one comparison but downregulated or unchanged in another (A) Mean values ± SD are represented with error bars. Statistical significance was determined by (A) two-way ANOVA, (C) Ordinary one-way Analysis of Variance (ANOVA).

### Transcriptomic profiling reveals reduced migration potential of Vδ1⁺ pTILsγδ and impaired effector potential in Vδ1⁺ mTILs_γδ_

To investigate whether the compartmental restriction of Vδ1⁺ γδT cells during metastatic progression is reflected in additional features of tissue-resident repertoires, transcriptomic analyses were performed using a publicly available single-cell RNA sequencing (scRNA-seq) dataset of colorectal cancer with liver metastasis (CRLM) samples. This dataset, hereafter referred to as CRLM (GSE164522), includes primary tumor-infiltrating lymphocytes (pTILs; n=7), lamina propria lymphocytes (LPLs; n = 8), liver metastasis-derived TILs (mTILs; n=7), and adjacent normal liver tissue samples (Liver_γδ_ ; n=9) ^30^. CD3⁺ TRDC⁺ CD8B⁻ γδT cells were computationally extracted based on gene expression and cell annotations, and differential gene expression analysis was conducted. Additionally, a patient-specific scRNA-seq dataset was generated from γδT cells of a CRC patient with liver metastasis (PAT39), and comparative analysis across both datasets was performed (Figure 6D). A comparison between pTILs_γδ_ and mTILs_γδ_ revealed significant upregulation of RGCC and TOB1 in mTILs_γδ_. These genes are associated with TGF-β–mediated suppression of T cell activation and cytotoxicity, as well as with promoting cellular quiescence^31,32^. Furthermore, KLRB1 was significantly upregulated in mTILs_γδ_, suggesting reduced TCR-mediated stimulation compared to pTILs ^33,34^. In contrast, pTILs exhibited significantly higher expression of genes associated with cytotoxicity and T cells survival, such as GZMB and TNFRSF9 (encoding 4-1BB), as well as activating receptors KLRC2 and KLRC3^35,36^. Interestingly, pTILs_γδ_ also expressed high levels of RGS1, a gene associated with reduced T cell trafficking^37,38^, potentially explaining their retention within the tumor microenvironment. Additionally, mTILs_γδ_ displayed high expression of GZMK along with lower GZMB, suggesting incomplete activation, or an early exhausted phenotype^39^. However, TIGIT, an exhaustion marker, was found at higher levels in pTILs_γδ_, while PDCD1 (PD-1) was not detected in either dataset. To validate transcriptomic observations at the protein level, GZMB and GZMK expression was assessed by flow cytometry in *ex vivo*-expanded Vδ1^+^ PBLs_γδ_, LPLs_γδ_, pTILs_γδ_, mTILs_γδ_. High levels of GZMB were confirmed in Vδ1^+^ pTILs_γδ_ from both tested patients. In contrast, high GZMK expression was detected only in the Vδ1^+^ mTILs_γδ_ population of one patient (Figure 6E). In conclusion, these data underscore the pivotal role of tissue origin in shaping the phenotype and function of γδT cells. Transcriptomic analyses revealed that highly tumor-reactive γδT cells within pTILs display enhanced cytotoxic signatures alongside features of reduced migratory capacity. In contrast, γδT cells from mTILs exhibit transcriptional profiles, indicative of impaired effector function and limited tumor reactivity.

### pTILsγδ, but not mTILsγδ, are protective in terms of overall survival

Our data suggest that pTILs_γδ_ may be clinically more relevant for tumor protection than mTILs_γδ_. To investigate this, we analyzed the prognostic impact of the frequency of pTILs_γδ_ and mTILs_γδ_ in primary tumors and matched liver metastases from a second independent cohort of 69 MSS patients with UICC Stage IV CRC, and a documented clinical history (Table 1, Cohort2). Tissue samples were analyzed by immunohistochemistry, focusing on the infiltrative margins of both primary tumors and liver metastases. Quantification of pTILs_γδ_ and mTILs_γδ_ was correlated with overall survival. Patients with high infiltration of pTILs_γδ_ had significantly improved overall survival compared to those with low or absent pTILs_γδ_ infiltration (P < 0.05; Figure 6F). In contrast, mTILs_γδ_ levels showed no association with overall survival (Figure 6G). In conclusion, consistent with their lack of tumor reactivity *in vitro*, mTILs_γδ_ do not appear to influence CRC progression. In contrast, the presence of pTILsγδ in primary MSS CRC is associated with a protective effect, and improved patient prognosis.

## Discussion

Understanding the behavior of γδT cells in solid tumors has been hindered by their rarity across tissue compartments, and the absence of integrative tools capable of linking functional screening, biomimetic migration assays, γδTCR repertoire analysis, and transcriptomic profiling. To address this major limitation and to map γδT cell states across different compartments in CRC patients, we developed a γδT-omics platform capable of linking different levels of characterization. Specifically, our functional γδT-omics platform initially enabled the isolation and expansion of γδT cell subsets amenable to in-depth functional and molecular characterization. This approach resulted in the functional characterization of γδT cell subsets from multiple expanded compartments of 12 MSS CRC patients, 7 healthy donors, and approximately 9000 individual γδT cell clones derived from PBLs, LPLs, pTILs, and mTILs. In a second phase, we contextualized these findings through γδTCR repertoire and transcriptomic analyses. These data were integrated with publicly available single-cell datasets^30^ to map compartment-specific γδT cell states in CRC, and to place the functional profiles of our expanded clones within a broader spatial and transcriptional context. Although the outgrowing cells analyzed might represent a subset of the broader tissue-resident γδT cell repertoire, and are inherently biased toward subsets capable of proliferating under *ex vivo* stimulation, we identified both highly tumor-reactive and poorly responsive clones. These findings suggest that our γδT-omics platform based on initial functional screens captures a physiologically relevant spectrum of γδT cell states, rather than being skewed exclusively toward CRC-reactive populations.

Our strategy revealed a striking compartmentalization of γδT cell function, migration, and clonality. Tumor-reactive Vδ1⁺ γδT cells were consistently enriched and clonally expanded in primary tumors, and derived predominantly from LPLs, as shown by shared γδTCR repertoires and a common CD69⁺CD103⁺ residency phenotype. The observed enrichment of tumor-reactive Vδ1^+^ γδTILs might also partly contribute to the lower tumor size associated with an increase of γδT cell infiltration, an upregulation of cytotoxic genes (e.g., GZMB, IFNG) in pTILs_γδ_ as well as their increased tumor-reactivity also against autologous targets as assessed by functional assays. These findings align with recent work highlighting tissue-imprinted Vδ1⁺ subsets as key cytotoxic players in epithelial tumors^10–13^. Transcriptomic analyses revealed that Vδ1⁺ LPLs_γδ_ and pTILs_γδ_ upregulate chemokine receptors (CCR5, CXCR3, CXCR4 and CXCR6) and ligands (CCL3, CCL4, CCL5) as also reported by others^18^, while we observed also a simultaneously downregulating of trafficking-related genes like RGS1 in pTILs_γδ_ when compared to LPLs_γδ_, a feature consistent with functional entrapment of Vδ1^+^ γδT cells upon tumor infiltration^11,37,38^. In line with this observation, our 3D migration assays using a trans-well system suggests that Vδ1⁺ LPLs_γδ_ migrate more efficiently than their tumor-infiltrated counterparts. This process initially depends on the CXCL16-CXCR6 axis and, upon cognate recognition of tumor cells via the γδTCR, is reinforced by the secretion of CCL3, CCL4, and CCL5 by tumor-infiltrating Vδ1⁺ γδT cells, thereby attracting CCR5⁺ Vδ1⁺ LPLs_γδ_. CCR5 was, however, not only expressed at a higher level at the cell membrane of expanded Vδ1⁺ LPLs_γδ_, but also downregulated once Vδ1⁺ LPLs_γδ_ came into contact with tumor cells, implying that CCR5 plays a role as a gatekeeper of Vδ1⁺ LPLs_γδ_ recruitment and on their subsequential entrapment in the tumor. These data imply that improved tumor trafficking via CCR5 engineering^40^ might be an interesting strategy for γδT cell-based therapies, as overexpression usually also avoids down-regulation.

Our study also sheds light on the functional divergence of Vδ1⁺ γδT cells between primary and metastatic sites. In contrast to the cytotoxic, clonally expanded Vδ1⁺ TILs in primary tumors, Vδ1⁺ γδT cells in liver metastases exhibited diminished tumor reactivity and a distinct, non-overlapping γδTCR repertoire. Indeed, distinct tissue specific γδT cell repertoires have been described not only in mice but also in human skin, lungs and the uterus^41^. Vδ1^+^ mTILs_γδ_ upregulated suppressive genes such as TOB1 and RGCC^31,32^, which are known to be induced by TGF-β and are associated with quiescence^32^. Our finding challenges earlier suggestions that γδT cells might maintain similar functionality across primary tumor and metastatic sites and instead supports a model in which local cues and tissue origin imprint unique γδT cell phenotypes. Vδ1^+^ γδT cells derived from metastatic sites are therefore most likely neither strongly CRC-reactive^29^ nor an optimal source for TIL therapies, in contrast to previous suggestions. Accordingly, our data showed that infiltration of pTILs_γδ_ in primary tumors is associated with a protective effect against CRC progression, whereas the presence of mTILs_γδ_ does not influence overall survival. These data also align with the observation that in MSS colorectal cancer, pathological responses to immunotherapy are occasionally seen in non-metastatic or primary tumors, but not in metastatic sites^42,43^.

A central question in γδT cell biology is whether tumor recognition is primarily mediated by innate co-receptors or by the γδTCR itself. Some studies suggested that NKG2D ^13,44,45^ or NKp30/44/46 ^46–48^ are the dominant mediators of anti-tumor responses in Vδ1⁺ γδT cells. Indeed, we observed high expression of NKG2D and NKp44 on Vδ1⁺ pTILs_γδ_ compared to HD Vδ1⁺ PBLs_γδ_ and confirmed persistent NKG2D expression following *ex vivo* expansion. However, functional blocking experiments showed no consistent correlation between co-receptor expression and tumor reactivity, with only a single patient demonstrating NKG2D-dependent cytotoxicity. To directly assess the impact of isolated γδTCR, we re-expressed Vδ1⁺ γδTCRs from tumor-reactive clones in CD4⁺ αβT cells (TEG format ^24,49,50^). These engineered cells preserved or even enhanced the original tumor reactivity, confirming that the γδTCR alone is frequently sufficient to confer CRC-specific recognition, and is most likely partially hampered through negative regulators expressed on primary Vδ1⁺ γδT cells. These findings reinforce the emerging view that γδTCRs not only encode tumor specificity, but also likely initiate the enhanced migration of Vδ1⁺ LPLs_γδ_ by inducing chemokine secretion upon antigen encounter. Moreover, γδTCRs can serve as scalable, antigen-independent tools for adoptive immunotherapy^51,52^. Such novel Vδ1^+^ γδTCRs will add to the arsenal of current cancer-specific γδTCRs, which are already in clinical testing ^53^ and have the potential to bypass the limitation of the lack of antigens in traditional CAR-T approaches. Moreover, engineering strategies may help address the natural migration constraints of tumor-reactive Vδ1⁺ γδT cells if paired with modifications to enhance their tumor trafficking, though ligand identification and safety profiling of novel Vδ1⁺ γδTCRs will be critical next steps to assure optimal clinical outcome.

Taken together, our results reveal a model in which tissue-resident Vδ1⁺ LPLs_γδ_ act as precursors to tumor-infiltrating γδT cells in MSS CRC. These cells acquire potent effector programs upon recruitment through CCR5 but become functionally entrapped due to RGS1-driven chemokine desensitization. The use of a comprehensive γδT-omics pipeline was essential for uncovering the spatially resolved interplay among trafficking, clonal adaptation, and effector function. It demonstrated that Vδ1⁺ mTILs_γδ_ are transcriptomically and functionally distinct from Vδ1⁺ LPLs_γδ_ and Vδ1⁺ pTILs_γδ_, and, unlike Vδ1⁺ pTILs_γδ_, do not confer protection against cancer progression. Beyond characterizing new γδTCRs for therapeutic development, our study provides a platform for understanding γδT cell behavior in other tissue-specific tumor contexts, opening new avenues for rational immunotherapy design, and for the identification of new potential therapy leads for solid cancer.

## Materials and Methods

### Patient material

In this study, we enrolled Colorectal Cancer (CRC) diagnosed patients who were scheduled for surgical resection at either the Beatrix Hospital Gorinchem, or the University Medical Centre Utrecht. The study was evaluated by the Biobank Research Ethics Committee (TCBio, 21-042) and conducted in accordance with good clinical practice guidelines and the principles of the Declaration of Helsinki. The selection of patients was based on microsatellite stability (MSS) and tumor location. All participants provided informed consent to the Utrecht Platform for Organoid Technology (UPORT), and the Prospective National CRC cohort (PLCRC) under TIL-studie (NL47888.041.14, METC 12/510). UPORT provides a logistical infrastructure for rapid and standardized acquisition of human tissues, based on approved protocols for patient inclusion www.uport.umcutrecht.nl. Upon informed consent and immediately after surgery, the UPORT platform provided primary tumoral, adjacent colon and CRC liver metastasis resections and the PLCRC pre-surgical blood. A total of 31 patients were enrolled in this study, and a summary of the patients with MSS CRC included in this study from this patient cohort 1is presented in Table 1.

A second patient cohort was used for immunohistochemical analysis of pTILs_γδ_ and mTILs_γδ_. FFPE-embedded tissues of primary tumors and matched liver metastasis from patients with MSS CRC were derived from the Comprehensive Cancer Center Freiburg Biobank of the University Medical Center, Freiburg. The patients underwent tumor surgery at the Medical Center Freiburg from 2011 – 2020. The study was conducted in accordance with good clinical practice guidelines and the principles of the Declaration of Helsinki. The study was approved by the local ethics committee of the University of Freiburg (ethics approval: 21-1162). A summary of the patients with MSS CRC included in this study from this second patient cohort 2 is presented in Table 1.

### Processing of blood

Sanquin Blood Bank (The Netherlands) provided buffy coat of healthy donors, while CRC patient pre-surgical blood was provided by PLCRC as mentioned above. Peripheral blood mononuclear cells (PBMCs) were isolated by density gradient centrifugation by the use of Ficoll-Paque (GE Healthcare). After of the isolation, the cells were cryopreserved or expanded by Rapid Expansion Protocol as described in the T cell cultures section.

#### Tissue collection and processing

Upon patient tissue collection, samples were immediately transferred to sterile advanced Dulbecco’s Modified Eagle Medium/F12 (Gibco), supplemented with 1× GlutaMAX Supplement (100X Gibco), 100 U/ml penicillin-streptomycin (Gibco), 10 mM HEPES (Gibco). Subsequently, the dimensions (height x width x length) of the biopsies were measured, and one-third of the tissue sample was used for organoid generation, leaving two-thirds of the sample for lymphocyte isolation. Biopsies were kept at at 4°C in Hank’s Balanced Salt Solution (HBSS, Gibco) containing 100 µg/mL Primocin (InVivoGen / Bio-Connect) (=HBBS^1^) during processing for lymphocyte isolation.

### Isolation of mononuclear cells

HBSS supplemented with 100µg/ml primocin was used to wash the biopsies and remove any excess debris and blood before mincing of the biopsies into small pieces and enzymatic digestion. The fragments were transferred to pre-digestion buffer consisting of HBBS^1^ containing 1mM DTT (Sigma D0632-5g) and 1mM EDTA, Fragments were continuously agitated during a 15 minutes incubation at 37°C before centrifugation at 1500 RPM for 5 minutes at 4°C and removal of the supernatant. This agitated incubation with pre-digestion buffer was repeated for a second cycle. Next, cells were incubated for 30min with gentle agitation at 37°C with digestion buffer consisting of HBBS^1^ supplemented with 0.05-0.375 units/ml DNase I (Invitrogen) and 2 mg/mL collagenase D (Roche/Merck). After digestion, the cell suspension was washed with HBBS and filtered using a 100 µm cell strainer (Falcon). Cell concentration and viability were determined by manual counting using a hemocytometer using both Turk’s solution and trypan blue. Finally, the lymphocytes cell suspension was analyzed by flow cytometry and cryopreserved. The count of live mononuclear cells, isolated from the different tissues, was normalized to the volume of the source and the immune cells density in the different compartments was determined.

### Organoid generation and culture

33% of the fragments of both the healthy and tumor tissue were used for the generation of organoid, according to the protocol of Cayetano Pleguezuelos-Manzano et al.^54^ and following the media instruction reported in Sato et al^55^. Samples were kept in media Advanced-DMEM/F-12+++ consisting in Advanced-DMEM/F-12 (Gibco) supplemented with 10mM Hepes (100x Gibco), 2mM glutamax (100x, Gibco) and 100u/ml pen/strep (Gibco). Fragments were digested for 60 min at 37°C in 5 mL Advanced-DMEM/F-12+++ containing 0.1mg/ml Liberase (Merck) and 10µM Y-27632 dihydrochloride (Tocris/Bio-Techne). The cell suspensions were washed in this Advanced-DMEM/F-12+++ three times before being resuspended in resuspended in Cultrex Basement Membrane Extract (BME, Bio-Techne) and seeded in three 3-dimensional droplets in a pre-warmed culture plate. This plate was inverted and incubated at 37°C for 20-60 30 min for the droplets to solidify. Next, expansion medium was added to each culture well consisting of Advanced-DMEM/F-12+++ supplemented with 50% 3dGRO™ R-Spondin-1 Conditioned Media Supplement (Merck; SCM104 ), 2% (v/v) Noggin conditioned medium (IPA), 1x B-27 Supplement (50x, Life Technologies), 1.25 mM N-acetylcysteine (Sigma-Aldrich), 0.5 µM ALK5 inhibitor A83-01 (Tocris/Bio-Techne), 50ng/mL recombinant human EGF (Peprotech), 50ng/ml recombinant human FGF2 basic (PeproTech), 100ng/ml recombinant human IGF-1 (Biolegend) and 0.1mg/ml primocin (InvivoGen/Bio-Connect). During the first 2 days of culturing, 10 µM Y-27632 (Tocris/Bio-Techne) was added to the cultures.1.25 mM N-acetylcysteine (Sigma-Aldrich), 10 µM p38 inhibitor SB202190 (Sigma-Aldrich), 0.5 µM ALK5 inhibitor A83-01 (Tocris/Bio-Techne), 10 mM nicotinamide (Sigma-Aldrich), 1 µM prostaglandin E2 (Tocris/Bio-Techne), 50 ng/mL epidermal growth factor (Peprotech), 2% (v/v) Noggin conditioned medium (U-Protein Express), 20% (v/v) Rspo1-CM (R&D Systems) and 1x B-27 Supplement (50x, Life Technologies). In the case of healthy colon organoid generation, expansion medium was supplemented with 0.5 nM Wnt surrogate (U-Protein Express Gibco). One dayTwo days post seeding of the droplets, expansion medium was refreshed with expansion medium w/o Y-27632. After a seven day incubation, organoids were passaged by refreshing both the expansion medium and BME according to the protocol published in Current Protocols in Immunology^54^.

### Flow cytometry

Isolated lymphocytes were characterized by staining with the following antibodies: panαβTCR-APC (IP26, eBioscience), panγδTCR-PE-Cy7 (IMMU510, Beckman Coulter), CD8α-PerCP-Cy5.5 (RPA-T8, BioLegend), CD8α-BV605 (RPA-T8, BioLegend), Vδ1-PE (REA173, Milteny), Vδ2-FITC (B6, BioLegend), CD3-PB (UCHT1, BD), DNAM-1 BV510 (11A8, BioLegend), CD45RA-APC-Cy7 (HI100, Sony Biotechnology), NKG2A-PE-Cy5 (S19004C, BioLegend), NKG2D-BV650 (1D11, BD), NKG2C-AF700 (134591, R&D), NKp46-PerCP-Cy5.5 (9E2, ThermoFisher), NKp44-BV786 (p44-8, BD), NKp46-PerCP-Cy5.5 (9E2, ThermoFisher), NKp30-BV711 (p30-15, BD), CD27-BV785 (O323, BioLegend), CD69-PE-Cy5 (FN50, BioLegend), CD56-APC-eF780 (NKH-1, Beckman Coulter), CD103-BV605 (Ber-ACT8, BioLegend), PD-1-BV711 (EH12.2H7, BioLegend), GZMK-FITC (370507, Biolegend), GZMB-APC (372204, Biolegend), CCR5-PE (313707, Biolegend), CXCR2-APC (320710, Biolegend), CXCR3-FITC (353704, Biolegend), CXCR4-BV605 (306521, Biolegend), CXCR6-PerCP5.5 (356009, Biolegend) and LAIR-1-BV510 (744558, BD).

Extracellular stainings were performed by diluting the required antibodies mix in FACS buffer, and incubating the cells at room temperature for 30 minutes in the dark. For the intracellular staining the BD Cytofix/Cytoperm (10482735, BD) was utilized according to the manufacturer’s instructions. After staining, all samples were analyzed on a BD LSRFortessa or BD Canto using FACSDiva or FlowJo software (BD Biosciences).

### T cell cultures

An *in vitro* Rapid Expansion Protocol (REP) was utilized in order to expand and culture primary patient derived T-cells and engineered αβT cells, following the 14-day protocol^23^. Primary T-cells were cultured in a culturing media named 10%huRPMI and composed of (RPMI-1640 medium (Gibco), 100 µg/mL primocin (ant-pm-2, Bio-connect), 100U/ml penicillin/streptomycin (100 U/mLGibco), 10% human serum (Sanquin) and 50µM beta-mercaptoethanol (Life Technologies). Engineered αβT cells were instead cultured in 2.5%huRPMI (RPMI-1640 medium supplemented with 100U/ml penicillin/streptomycin (100 U/mLGIBCO), 2.5% human serum (Sanquin), and 50µM beta-mercaptoethanol (0.05M)). Partial medium change was performed every 2-3 days of cultures, in the huRPMI media with additional IL-2 (50U/mL). For primary T-cells 5ng/mL of IL-15 were was also added. All T cell cultures were kept at 37°C and 5% CO₂ and underwent multiple REP cycles to obtain sufficient T cell numbers for the subsequent TCR sequencing, functional assays and cell sorting.

### FACS cell sorting

T cells were stained using the following antibody mix before FACS cell sorting: panαβTCR-APC (IP26, eBioscience), CD3-PB (UCHT1, BD), CD8α-PerCP-Cy5.5 (RPA-T8, BioLegend), panγδTCR-PE-Cy7 (IMMU510, Beckman Coulter), Vδ1-PE (REA173, Milteny), Vδ2-FITC (B6, BioLegend) by incubation for 30 minutes at 4°C. Cell sorting was performed on the BD FACSAria II Cell sorter, resulting in the isolation of the following subsets; αβTCR+, Vδ2+, Vδ1+, and Vδ1-/2-. Single T cells were placed in 96-well plates containing 50 µL of 10% huRPMI and REP mix. Bulk T cells were sorted in 5 mL tubes also containing 10% huRPMI and REP mix.

### Cell line cultures

The suspension tumor cell lines Daudi, RPMI-8226, ML-1, and K562 were cultured in RPMI-1640 medium containing 100U/ml penicillin/streptomycin (Gibco) and 10% fetal calf serum (FCS) The adherent, solid tumor cell lines SCC9, Caco-2, HT-29, RA, and U87 were cultured in DMEM+GlutaMAX with the addition of 100U/ml penicillin/streptomycin (Gibco) and 10% FCS (Bodinco). Detachment of the adherent cell lines was achieved by utilizing a 0.25% trypsin-EDTA solution in PBS. These cell lines were passaged two times a week to a concentration 0.2 million cells/ml and incubated at 37°C with 5% CO2. For functional assays, cells from passage 4 up to passage 25 after thawing were used.

### IFNγ ELISpots

In order determine tumor-reactivity, IFNγ ELISpot assays were used on both T cell clones, T cell bulks and engineered human T cells after co-cultures, with different tumor cell lines and the generated CRC organoids. To this end, ELISpot plates (Millipore Multiscreen™ plates) were coated with 5 µg/mL of anti-IFNγ mAb (1-D1K, Mabtech). For testing of both primary γδT cell populations and γδTCR transduced cells, a 1:3 ratio was used, comprised of 15,000 effector cells and 50,000 target cells. In case of blocking experiments, T-cells were pre-incubated for 2h 37C with 10µg/mL, the neutralizing antibody NKG2D (MAB139, R&D systems) or IgG isotype as a negative control (110-HG-100, R&D system).

As a positive control, T cells were stimulated with 1 µg/mL ionomycin (Sigma) and 20 ng/mL Phorbol 12-Myristate 13-Acetate (PMA) (Sigma-Aldrich), while the negative control wells only contained T cells without target cells. 30 µg/mL. Following the overnight co-culture at 37°C, 5% CO2, cell plates were washed, and cells were removed. IFN-γ-producing T cells were identified utilizing 1 µg/mL of biotinylated anti-IFNγ antibody (clone 7-B6-1; Mabtech) in PBS containing 0.5% FCS, followed by the administration of streptavidin-HRP (Mabtech, diluted 1:500) and TMB substrate solution (Sanquin). The total count of IFNγ positive spots was determined by the automated ELISpot reader ELISPOT Analysis Software (Aelvis) according to the following parameters: minimum intensity 2, minimum circularity 30, minimum size 2 and maximum size 500. To calculate the frequency of tumor reactive clones, a tumor-reactive clone was defined by ≥ 30 IFNγ spots released, compared to T cell-only controls against at least one tumor cell line.

### Luciferase based killing assay

In case of blocking experiments, T-cells were previously blocked for 2h at 37°C with 10µg/ml of NKG2D (MAB139, R&D systems) and NKp46 (MAB1850, R&D systems) neutralizing antibodies. Primary Vδ1 or αβ, transduced with either the Vδ1 or Vδ2 TCRs, were incubated with the different tumor cell lines SCC9, HT29, or RPMI-8226 cells containing luciferase GFP at different E:T ratios. After a 24 hour co-culture, 125 µg/mL of Luciferine was added after which luminescence signal was measured using the Softmax Pro machine. A tumor target only condition was used to normalize the data.

### γδTCR sequencing of expanded vd2-clones

The RNeasy Micro Kit (Qiagen) was used to isolate RNA of Vδ2-clones (500,000 cells) following the manufacturer’s guidelines. Next, the RNA was converted to cDNA utilizing the the iScriptTM cDNA Synthesis Kit (Bio-Rad, #1708890) according to the manufacturer’s protocol. The synthesized cDNA was amplified by PCR using the following reverse primers for the constant region: TRDCRev: TTCACCAGACAAGCGACA, TRGCRev: GGGGAAACATCTGCATCA. A mixture of forward primers were a used specific for the γ chain and δ chain; TRGV2-5Fw: CTGCCAGTCAGAAATCTTCC, TRGV8Fw: GCTGTTGGCTCTAGCTCTG, TRGV9Fw: TATGGTGCAGGTCACCTAGAG, TRDV1Fw: AGTGTGGCCCAGAAGGTTAC, TRDV2Fw: TCTCTTCTGGGCAGGAGTC, TRDV3Fw: GCACGCTGTGTGACAAAGTA, TRDV4Fw: CAGCTTCACTGTGGCTAGG, TRDV5Fw: GGGCATCAGTGCTGATTC, TRDV6Fw: GCTTCAACTATGCTGGGTGAG, TRDV7Fw: CTTCTACTGAGCTGGGTGAG, TRDV8Fw: CTTGTGATCTCCACCTGTCTTG.

Amplification was performed on a SimpliAmp Thermal Cycler with the Q5 High-Fidelity DNA Polymerase. PCR cycling was performed according to the following settings; denaturation at 95°C for 2 minutes, followed by 35 cycles consisting of 98°C for 20 seconds, 57.6°C for 20 seconds, and 72°C for 45 seconds, with the final extension of 7 minutes at 72°C. PCR products were run on a 1% agarose gel and subsequently purified following the manufacturer’s protocol of the Gel Clean-up kit (Macherey-Nagel). The purity and concentration of the DNA were determined using the NanoDrop spectrophotometer (Thermo Fisher Scientific). Macrogen conducted Sanger sequencing on the samples by utilizing the primer pairs used in the PCR amplification. TCR sequences were aligned in the IMGT-V-QUEST program (version 3.6.1) to analyze the sequencing FASTQ data.

### Cloning γδTCR sequences in pBullet plasmid

Overlap PCRs were used to reconstruct the matched γ chain and δ chain of the TCRs found in the monoclonal γδT cell populations, in separate retroviral expression vectors. Next, the amino acid sequences of the γ and δ chains were codon optimized by the Codon Optimization tool of Integrated DNA Technologies. For the overlap PCRs, the forward and reverse primers were designed to flank the CDR3 sequences for integration of both the γ or δ chain sequences into the expression vectors. The primer design was facilitated by the Clone Manager Professional 9 software (Sci Ed Software). In order to amplify the VDJ regions of the TCRs, a three-step approach was used: PCR reaction R1, R2 and R3. Part of the CDR3 region and variable domain were amplified in PCR R1 using the TP2F4 or TP1G1 TCR-specific reverse primer combined with the forward primers binding the template chains γ and δ TCRG115 clones: G115δFwd: CTG*CCATGG*AGCGGATCAGC, G115γFwd: G*CCATGG*TGTCCCTGCTG. For PCR R2, the remaining part of the TCR chains was amplified utilizing the 04TP4C4, 04TP2F4, 09TP1G1 or 14TP2C3 TCR-specific forward primer with the following constant reverse primers: G115δRev: ATGC*GGATCC*TCACAGG, G115γRev: TAGT*GGATCC*TCAGCTCTTCTC.

The R1 and R2 PCR products were run on a SimpliAmp Thermal Cycler (Thermofisher) with the following cycling settings: initial denaturation at 95°C for 2 minutes, followed by 30 cycles consisting of denaturation for 20 seconds at 95°C, annealing for 20 seconds at 59°C, extension for 40 seconds at 72°C and a final extension for 10 minutes at 72°C. The PCR products underwent agarose gel electrophoresis in a 1% agarose gel in TAE buffer and were subsequently purified by the Macherey-Nagel Gel Clean-Up Kit. In order to generate the TCR chains containing the TP2F4 or TP1G1 TCR-specific CDR3 regions, PCR products of the R1 and R2 reactions were combined and underwent the R3 PCR amplification with the G115γFwd/G115γRev and G115δFwd/G115δRev primer pairs. Next, the synthesized γ or δ chain sequences were purified and cloned into retroviral pBullet vectors using the NcoI and BamHI restriction sites included in the primers from R1 and R2. The TCRδ genes were ligated into the pBullet-IRES-puromycin vector, while the pBullet-IRES-neo vector was used to insert the TCRγ genes. Finally, sequences of the cloned TCR genes were verified by Macrogen using Sanger sequencing.

### Transduction of αβT cells

Retroviral particles were generated by overnight incubation of Phoenix Ampho cells at a cell density of 5 x 10⁵ cells per well in DMEM+GlutaMAX (Gibco) containing 10% FCS, and 1% penicillin (100 U/ml), and streptomycin (100µg/ml), in 6-well plates. The retroviral pBullet constructs containing the DNA of both the TCRδ and TCRγ chains of either the TP1G1 or TP2F4 TCRs were mixed with the packaging plasmids pHIT60 (containing gag-pol genes), and pCOLt-GALV (containing the ENV gene) in the presence of FuGENE HD (Promega). This transfection mix was administered to the Phoenix Ampho cells after a 20 minute incubation at room temperature. After an overnight incubation, the medium was replaced by 5% huRPMI consisting of RPMI-1640 medium, supplemented with 1% penicillin (100 U/mL), streptomycin (100 µg/mL), 5% Human Serum and 0.1% of beta mercaptoethanol (0.05M). Magnetic bead separation (Miltenyi Biotec) facilitated the isolation of CD3+ T cells from PBMCs according to the manufacturer’s protocol, after which T cells were cultured in 5% huRPMI. Next, administration of anti-CD3/CD28 antibodies (Dynabeads Human T-Activator CD3/CD28, Thermo Fisher Scientific), 25 IU/mL IL-15 and 1700 IU/mL IL-7 allowed for the activation of the isolated T cells. Upon activation, T cells were transduced via a two-hit transduction protocol. Viral supernatant of the Phoenix Ampho cells was collected at both 48 hours and 72 hours post-transfection, and filtered through a 0.45 µm syringe filter. During the first hit, viral supernatants supplemented with polybrene (8 μg/mL), 1700 IU/mL IL-7 and 25 IU/mL IL-15 were administered to the T cells, and incubated overnight at 37°C. The second hit was performed using fresh, viral supernatants containing 6 µg/mL Polybrene (Sigma-Aldrich), after which cells were rested at 37°C for 2 hours. Upon a 2-hour incubation, transduced T cells were cultured in 5% huRPMI containing 800 µg/mL G418 neomycin (Thermo Scientific), 1700 IU/mL IL-7, and 25 IU/mL IL-15 to select for transduced cells. After a five day incubation, T cell cultures were supplemented with a combination of 5 µg/mL puromycin (Merck), 25 IU/mL IL-15 and 1700 IU/mL IL-7. Seven days later, transduced T cells underwent magnetic bead depletion for the removal of transduced cells, and were subsequently put on the REP protocol.

### Trans-well migration assays

CACO2 tumor cells were stained with Vybrant DiO (Thermo Fisher) and seeded in basement membrane extract (reduced growth factors) (R&D Systems) as previously described in detail^56^, in the lower compartment of a 5.0 uM trans-well plate (Corning). After a four 4 day incubation, different γδT cell subsets, or TEGs, were administered to the upper compartment of the trans-well insert, together with 10 μM PAM in the case of the Vδ2+ TEG condition (Calbiochem). In case of chemokine receptors blocking experiments, primary T-cells and TEGs were pre-incubated for 2h at 37C, with either 100nM of the CCR5 inhibitor Maraviroc (HY-13004, Bio-connect), or 5µM of the neutralizing antibody of CXCR2 (MAB331, R&D system). Two days post T cell administration, BME was dissolved using cell recovery solution (Corning) according to the manufacturer’s protocol, in order to retrieve the T cells from both compartments of the model. T cells were quantified by FACS utilizing Flow-Count Fluorospheres (Beckman Coulter). Supernatant of the lower compartment at the end of migration was used to performed Luminex assay and measure levels of CCL3, CCL4, CCL5, CXCL5 and CXCL6.

### Secretome analyses

HT-29 cells were plated in quadruplicates at a density of 2.2×10^6^. After 48h, cell-conditioned media (CCM) of each of the replicates was collected separately. Fresh medium was added to the cells, collected again after 48h, and combined to the previously collected CCM. Samples were centrifuged at 1200 rpm for 5 minutes to remove cells and debris, and supernatants were transferred to fresh tubes. CCM were concentrated using 3kDa cutoff filters and mixed 1:1 with lysis buffer (10% SDS, 100mM TEAB, pH 7.5). Samples were reduced, alkylated, and processed using the S-Trap spin column digestion protocol. 16plex TMT labeling was performed using 12 channels, and samples were pre-fractioned on an Agilent-HPLC-system into 16 fractions. MS samples were measured on a Q Exactive Plus mass spectrometer using standard TMT parameters.MS files were analyzed using MaxQuant (parameters: Rep. Mass tol. 0.003, Min. reporter PIF 0.75, no variable modifications, Trypsin digest 2 missed cleavages, database: Mouse-EBI-reference, at least 1 unique peptide for quantification) and further processed for statistical analysis using Limma (for multiple group ’ANOVA’ and pairwise comparisons).

### Immunohistology and image analysis

FFPE embedded tissue from primary tumors and liver metastases from patients with colorectal cancer were sliced into 3 µm thick sections. For immunohistochemical staining, the slides were deparaffinized with Rotihistol (Carl Roth), rehydrated through a descending alcohol series, and washed with H₂O. Antigen retrieval was performed using pressure cooking (Retriever, Aptum Biologics) in 20 mM Citrate Buffer (pH 6.0, Zytomed Systems). To reduce endogenous peroxidase activity, the samples were incubated with 3 % H₂O₂ for 30 minutes. After blocking with 5 % goat serum (Sigma Aldrich) for 1 hour, the slides were incubated overnight at 4°C with the primary antibody (dilution 1: 500, TRDC/TCRδ (E2E9T) #55750, Cell Signaling Technology) in 1 % goat serum. Following primary antibody incubation, the slides were washed, and the primary antibodies were detected using anti-rabbit polymer (Zytomed Systems), according to the manufacturer’s instructions. DABplus (Zytomed Systems) was applied, followed by counterstaining with hematoxylin (Merck), dehydration in an ascending alcohol series, and Rotihistol and mounting with Rotihistokit (Roth). The slides were then digitized using the Zeiss Axio Scan Z.1 Microscope Slide Scanner (Zeiss). T cells were analyzed in the infiltrative margin of primary tumors and liver metastases. Five equal-sized ROI (region of interests) randomly dispered over the infiltrative margin were analyzed. pTILs_γδ_ or mTILs_γδ_ within the ROI were counted using QuPath^57^. Survival curves were estimated with the Kaplan-Meier method. Overall survival was analyzed by log-rank test, and results were expressed as hazard ratios (HR) with their 95% confidence intervals (95% CI). Score density of pTILs_γδ_ at the invasive margin was assessed as follows: 0, absent; 1+, sparsely (< 10 cell/mm^2^); 2+, moderate (10–20 cells/mm^2^); 3+, frequent (> 20 cells/mm^2^) and their density was evaluated as a dichotomous variable as high (2+, 3+) versus low (0, 1+) pTILs_γδ_ infiltration. Score density of mTILs_γδ_ at the invasive margin was assessed as follows: 0, absent; 1+, sparsely (< 5 cell/mm^2^); 2+, moderate (5–10 cells/mm^2^); 3+, frequent (> 10 cells/mm^2^) and their density was evaluated as a dichotomous variable as high (2+, 3+) versus low (0, 1+) mTILs_γδ_ infiltration.

### Bulk TCR-seq analysis

Fresh primary γδT cells were used to isolate RNA, generate cDNA and expand it in PCR with Merck primers(FW: 5’CTGTCAACTTCAAGAAAGCAGCGAAATC3’; RV:5’GTAGAATTCCTTCACCAGACAAG-3’) in order to amplify the CDR3 regions. CDR3 regions of TRD were amplified via gene-specific primers^51,58^. Libraries were sequenced. Annotation of FASTq files was performed with the MiXCR software^59^ using the *milab-human-tcr-dna-multiplex-cdr3* preset. Annotated read files were summarized using immunarch (https://immunarch.com/index.html).

### Single-cell RNA/TCR sequencing

Frozen patient samples processed and frozen on the day of surgery, were thawed and prepared for sorting following the methods described in the corresponding paragraph above. Sorting was performed per sample using BD FACSAria II Cell sorter, resulting in the isolation of the following subsets: CD3+αβTCR+ and CD3+γδTCR+ in a 1:1ratio. Cooling was maintained upon sorting, and cells were kept on ice during the whole procedure. Libraries for single-cell transcriptome sequencing and scTCR-seq were prepared using the Chromium Single-Cell 5′ Library GEM-X Gel Bead and Construction Kit and Chromium Single-Cell V(D)J Enrichment Kit (v3, 10x Genomics, CA, USA). Custom primers for the capture of γδTCR transcripts were produced as described by Tan et al^60^. The scRNA-seq and scTCR-seq libraries were sequenced on the Illumina Novaseq 6000 platform.

### Data processing of scRNA-seq libraries

Alignment to the reference genome (GRCh38) and generation of cell-gene matrices were performed using Cell Ranger v3.1 (10x Genomics). The resulting count matrix was analyzed in R software using Seurat v5.1 package^61^. Gene expression profiles were log(transformed) and scaled to 10,000 counts. We utilized principal component analysis (PCA) for linear and uniform manifold approximation and projection (UMAP) for non-linear dimensionality reduction, and single-cell embedding (https://arxiv.org/abs/1802.03426). Cells were clustered using FindClusters function from the Seurat package, and differentially expressed genes were identified using FindMarkers function from the Seurat package. For scTCR-seq analysis, Cell Ranger VDJ tools v3.1 were used to generate scTCR annotations, with GRCh38 as the alignment reference. Non-productive TCR sequences and duplicates were excluded. Additionally, scTCR-seq data were integrated into the scRNA-seq Seurat project metadata by matching cell barcodes.

### Analysis of public scRNA-seq datasets

For the pCRC dataset, we downloaded publicly available scRNA-seq data from the gene expression omnibus (GEO) database with the accession number of GSE178341^28^. Based on the original annotation, γδT cells from mismatch repair potent (MMRp) samples were extracted. For the CRLM dataset, we downloaded publicly available scRNA-seq from GEO with the accession number of GSE164522^30^, and extracted CD3^+^TRDC^+^CD8B^-^ γδT cells from the dataset. We analyzed both datasets using normal a Seurat V5.1 pipeline, and embedded cells in 2D using UMAP.

### Statistical analysis

Statistical analyses were performed using the GraphPad Prism software (version 9, GraphPad Inc.). For datasets containing independent samples, or those not meeting the assumption of normality, the Kruskal-Wallis test was used, followed by the Dunn’s post hoc, multiple comparisons test. Evaluation of differences among various groups was determined by a two-way analysis of variance (ANOVA). In cases where significant variance was observed, an additional one-way ANOVA was performed. Next to the ANOVA, Tukey’s or Šidák’s multiple comparisons tests were also applied. Statistical significance was determined by P < 0.05, with significance levels shown using the following symbols; *P < 0.05, **P < 0.01, ***P < 0.001, and ****P < 0.0001.

### Data and materials availability

Upon revision, scRNA-seq and scTCR-seq datasets will be deposited online in the GEO.

## Acknowledgements

We thank the staff of the Flow Core Facility at the UMC Utrecht for their kind assistance. Human tissues and other body materials, such as blood, were obtained with the help of the Utrecht Platform for Organoid Technology (UPORT) and the Prospective National CRC cohort (PLCRC). UPORT provides a logistical infrastructure for rapid and standardized acquisition of human tissues, based on approved protocols for patient inclusion www.uport.umcutrecht.nl. We thank the UMCU Pathology department for processing and delivering the samples. Illustrations were created by BioRender.com.

## Author contributions

Conceptualization: LG, MN, ZS, JK; Methodology: LG, MN, FK, PB, AM, ZS, JK; Formal analysis: LG, FK, PB; Investigation: all authors. Resources: OK, SM, JR. Writing-Original draft: LG, MN, ZS, JK. Writing editing: LG, MN, FK, PB, AM, DB, RK, SM, JR, ZS, JK. Writing revision: all authors. Supervision: ZS, JK.

## Supplementary Figures

**Supplementary Fig. 1.**
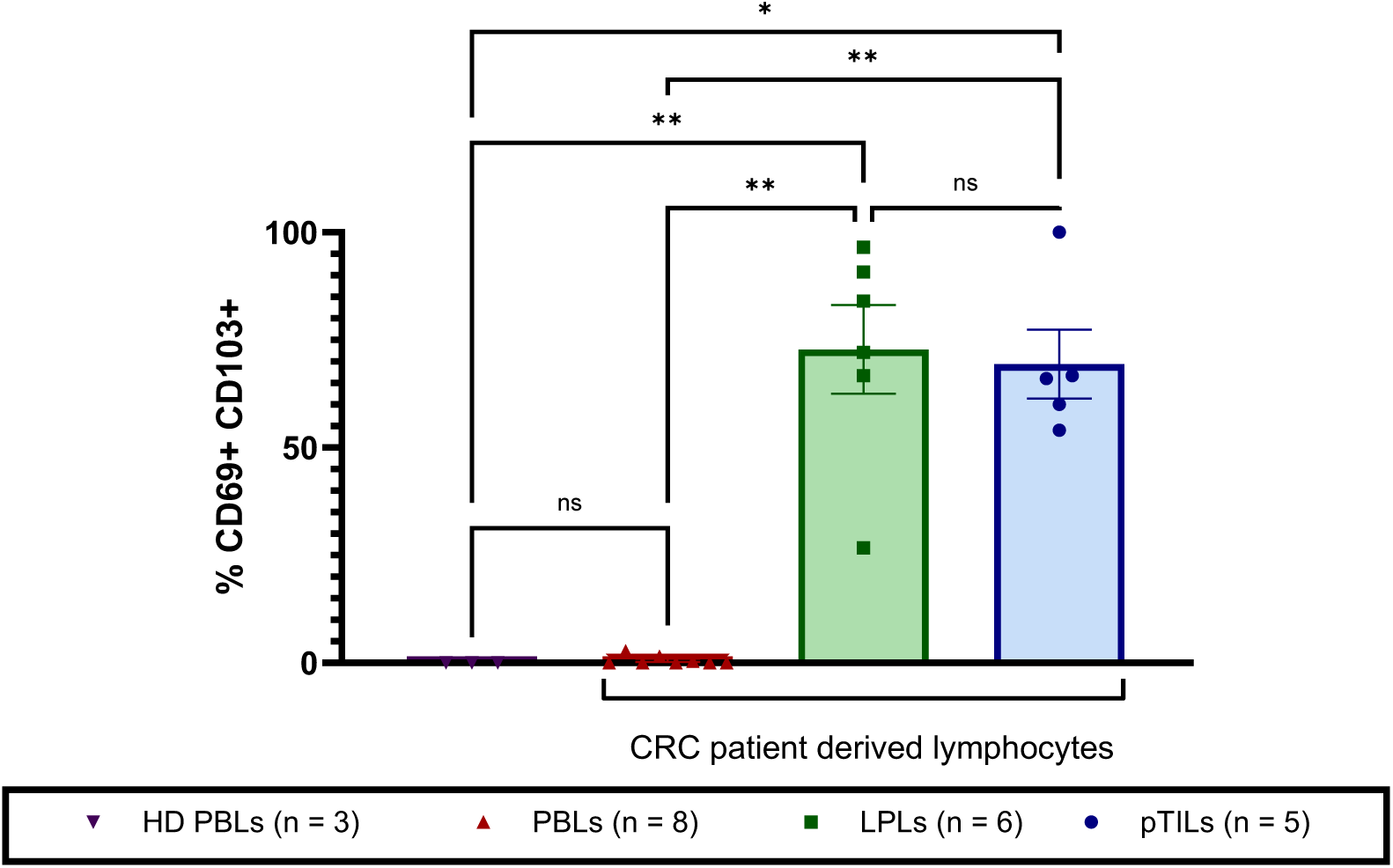
Tissue residency markers (CD69+CD103+) on Patient primary Tumor-infiltrating Lymphocytes (pTILs), Patient Lamina Propria Lymphocytes (LPLs), and both Patient and healthy donor (HD) Peripheral Blood Lymphocytes (PBLs). Individual data points indicate values from individual CRC patients or HDs. Not all patients had paired samples. Mean values ± SEM are represented with error bars. Statistical significance was determined by Ordinary one-way ANOVA.

**Supplementary Figure 2:**
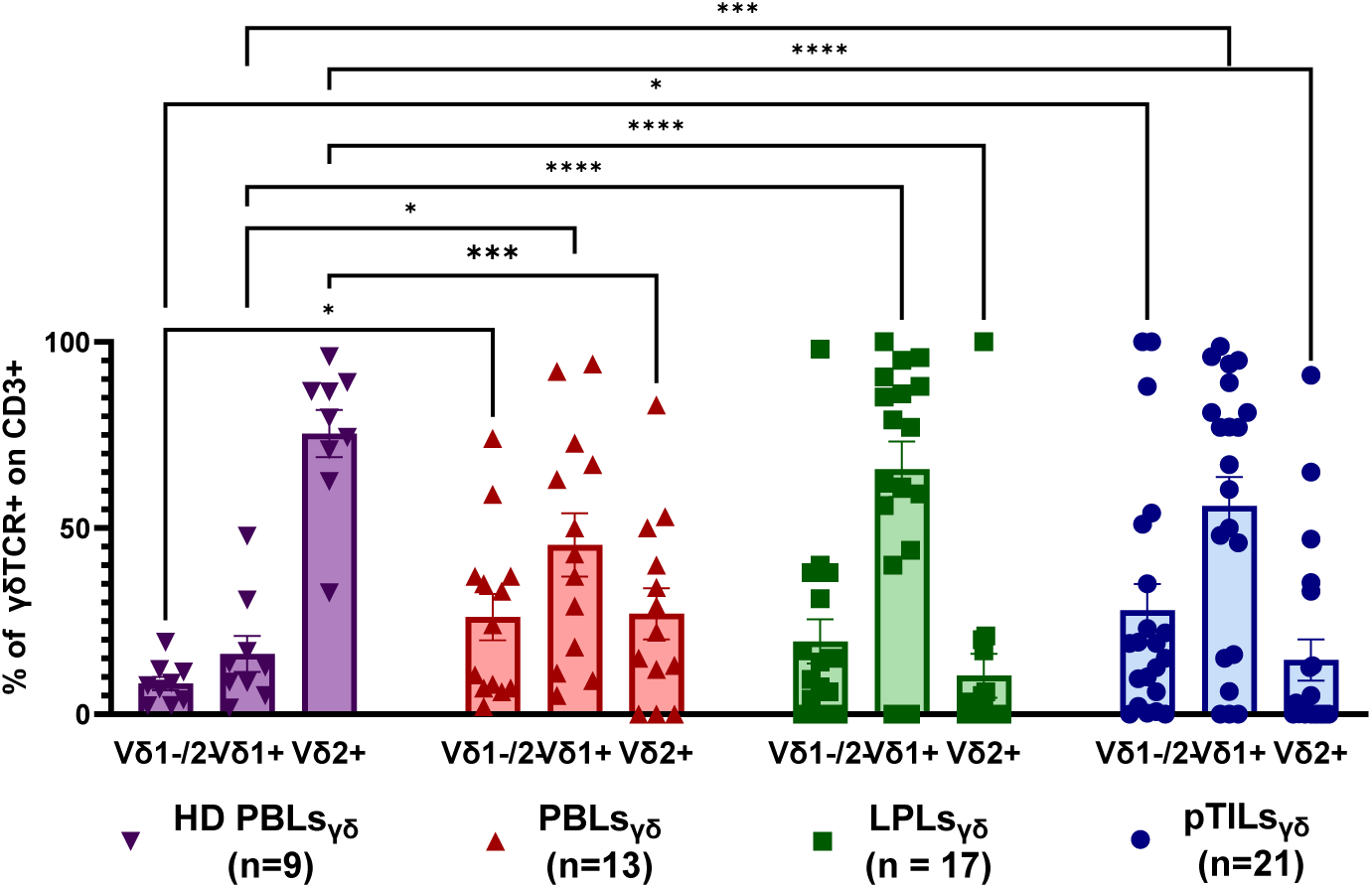
percentages of γδT cell subsets in the different patient compartments, PBLs_γδ_, LPLs_γδ_, pTILs_γδ_,_,_ and HD PBLs_γδ_ measured by flow cytometry in unexpanded γδT cells.

**Supplementary Fig. 3.**
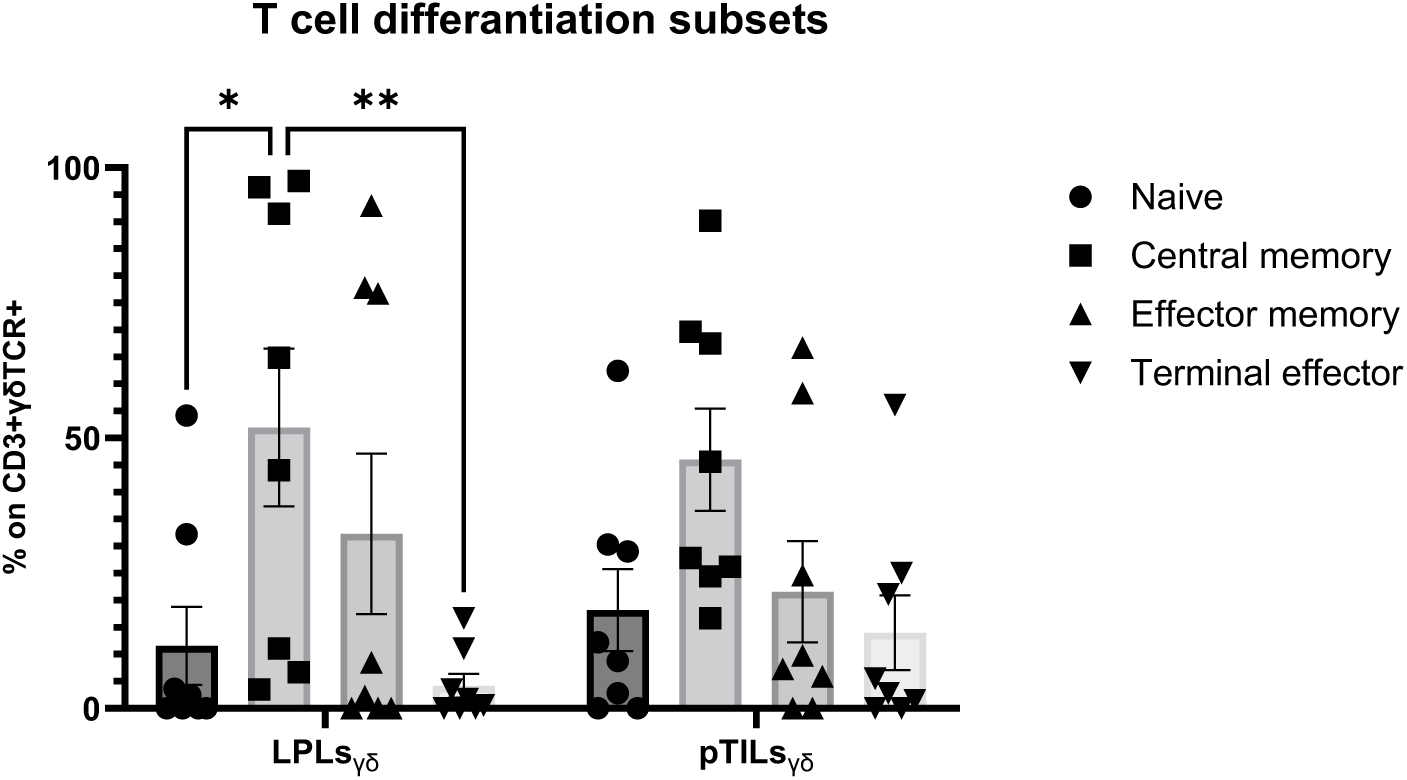
Cell differentiation of γδT cells derived from LPLs_γδ_ and pTILs_γδ_ based on CD45RA and CD27 expression. The majority of γδT cells derived from TILs_γδ_ and LPLs_γδ_ were central memory T cells. Not all patients had paired samples. Mean values ± SEM are represented with error bars. Statistical significance was determined by two-way ANOVA.

**Supplementary Fig.4.**
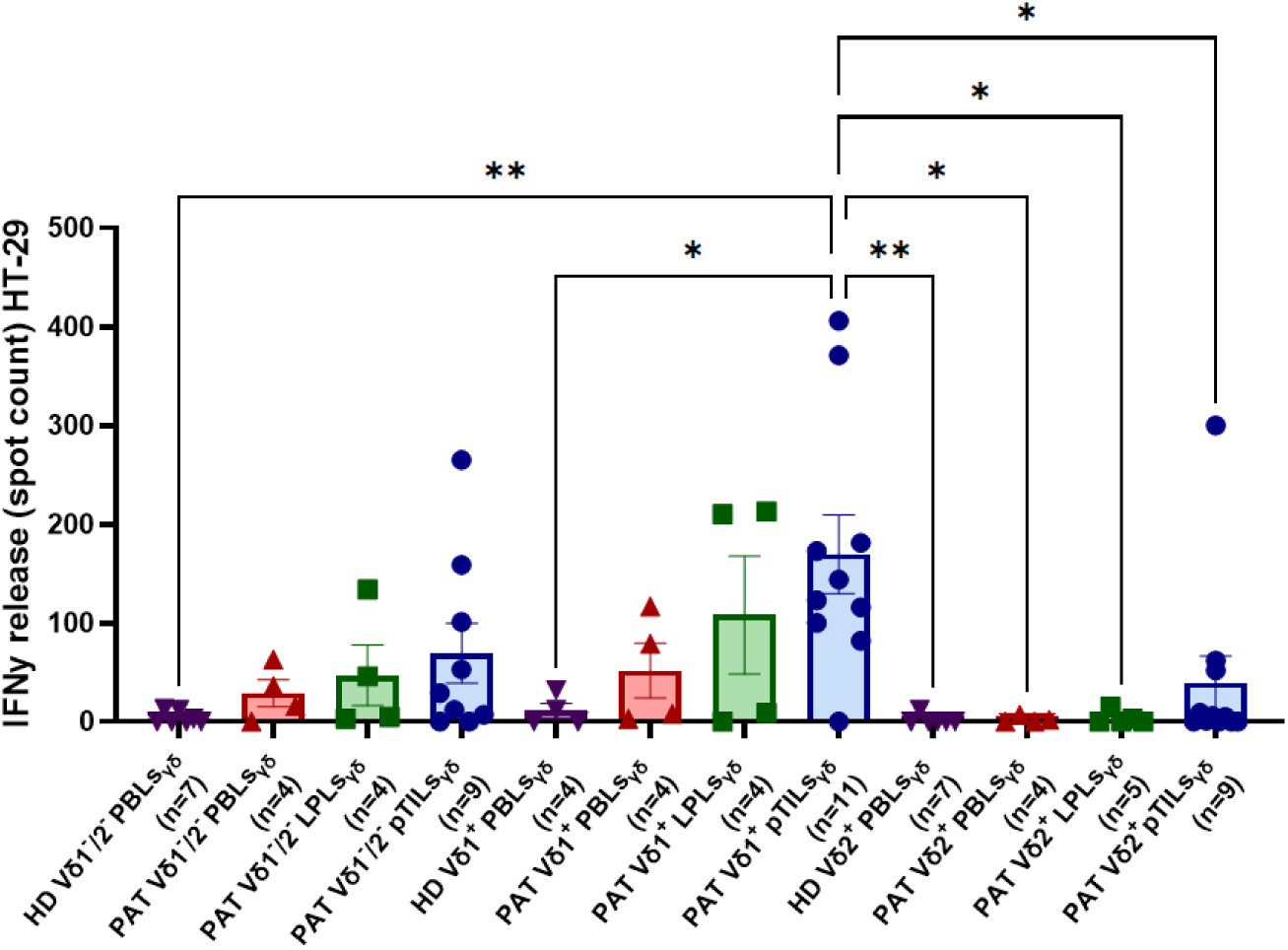
IFNγ release by either Vδ1^+^, Vδ2^+^, or Vδ1^-^/2^-^ expanded γδT cell bulks derived from the different sources in response to co-culture with HT-29 and Caco-2 cell lines. Tumor reactivity was assessed after a 24-hour co-culture of T cells with CRC cell lines at a 1:3 ratio, normalized to T cell only conditions, using an IFNγ ELISpot assay.

**Supplementary Fig. 5.**
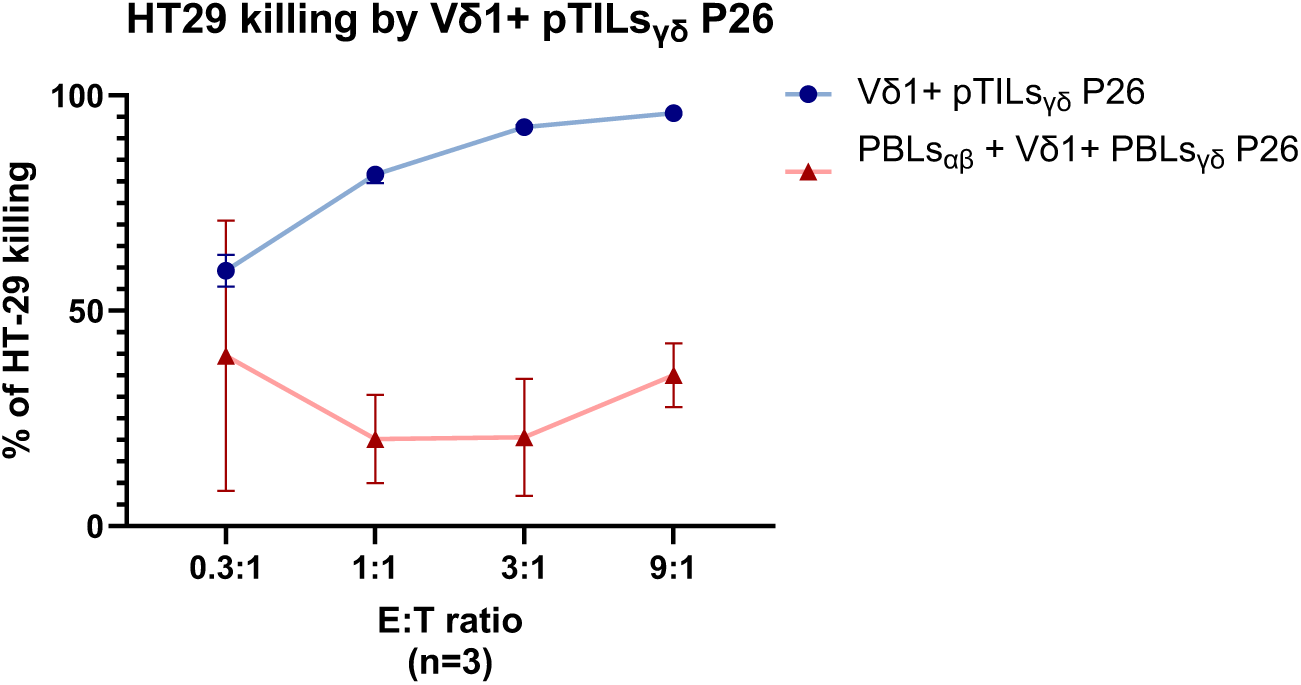
Luciferase based killing by the Vδ1+ pTILs_γδ_ and a combination of PBLs_αβ_ and Vδ1+ PBLs_γδ_ from the same patient, targeting the CRC tumor cell line HT-29 at multiple E:T ratios for 30h, normalized to the target only condition. n=3.

**Supplementary Fig. 6.**
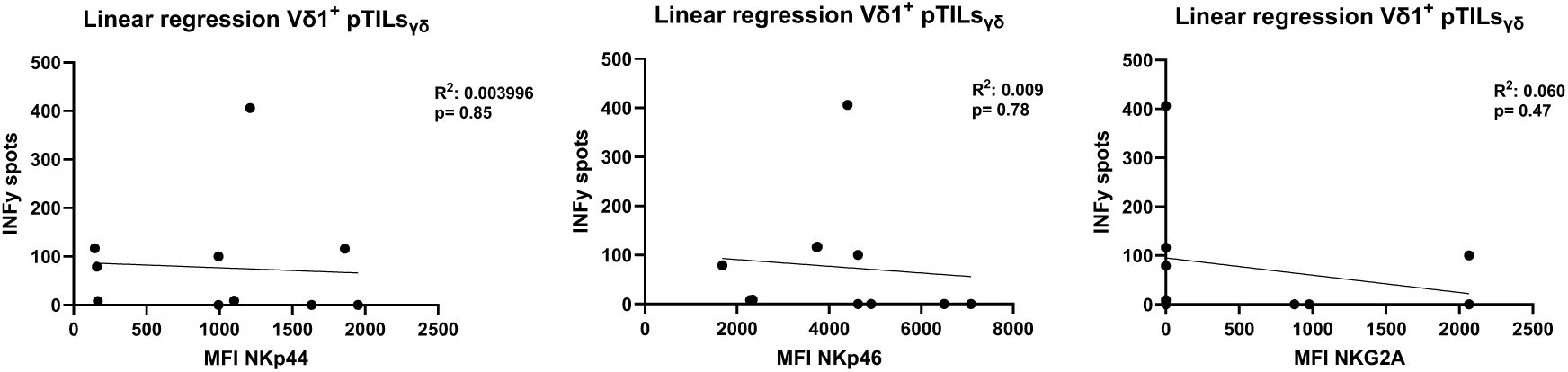
Linear regression between the NKp44, NKp46, NKG2A expression of the expanded Vδ1+ pTILs_γδ_ and IFNγ release against HT-29. Each data point represents paired measurements of IFNγ secretion and co-receptor expression from individual co-cultures. Correlation statistics were determined using simple linear regression analysis.

**Supplementary Fig. 7.**
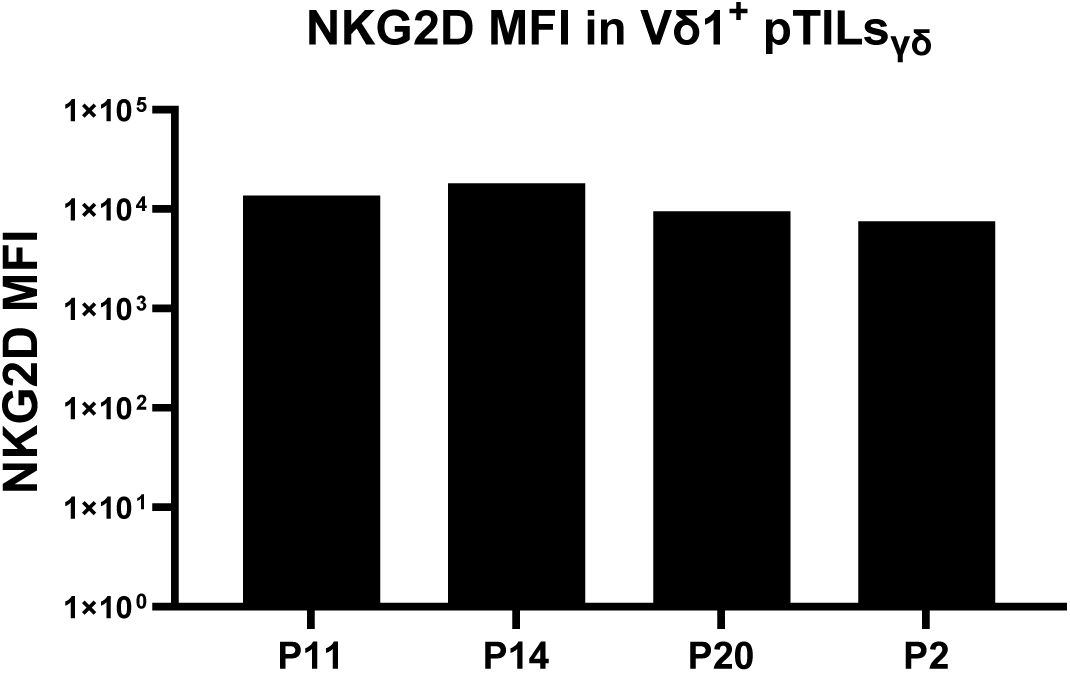
MFI of NKG2D surface expression on tumor reactive Vδ1+ pTILs_γδ_ derived from 5 patients measured by FACS (Fortessa BD) on the same day of functional tests.

**Supplementary Fig. 8.**
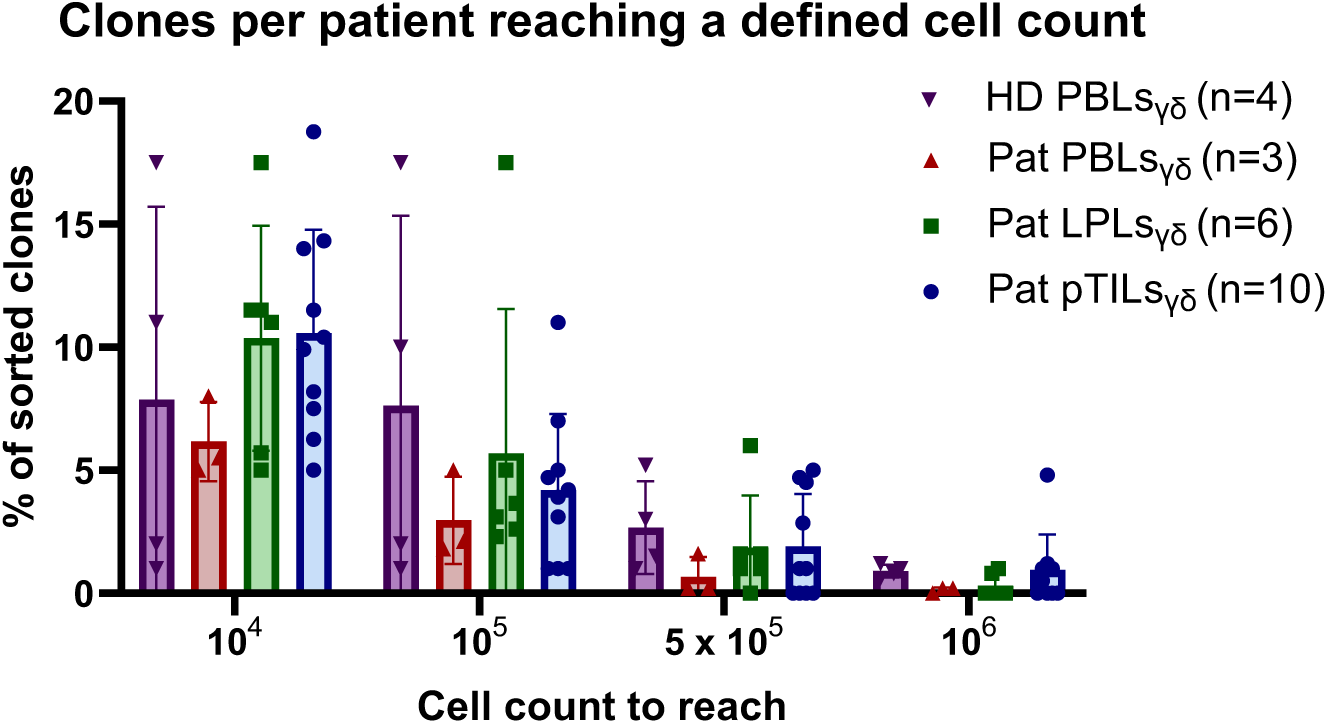
Percentage of single cell sorted T cell clones, isolated from either Pat pTILs, Pat LPLs, Pat PBMCs of CRC Patients, or from PBMCs of HD, that were able to be expanded up to a defined cell count (10^4^, 10^5^, 5×10^5^, 10^6^). Every dot represents a patient. Mean values ± SD are represented with error bars. Statistical significance was determined by two-way ANOVA.

**Supplementary Fig. 9.**
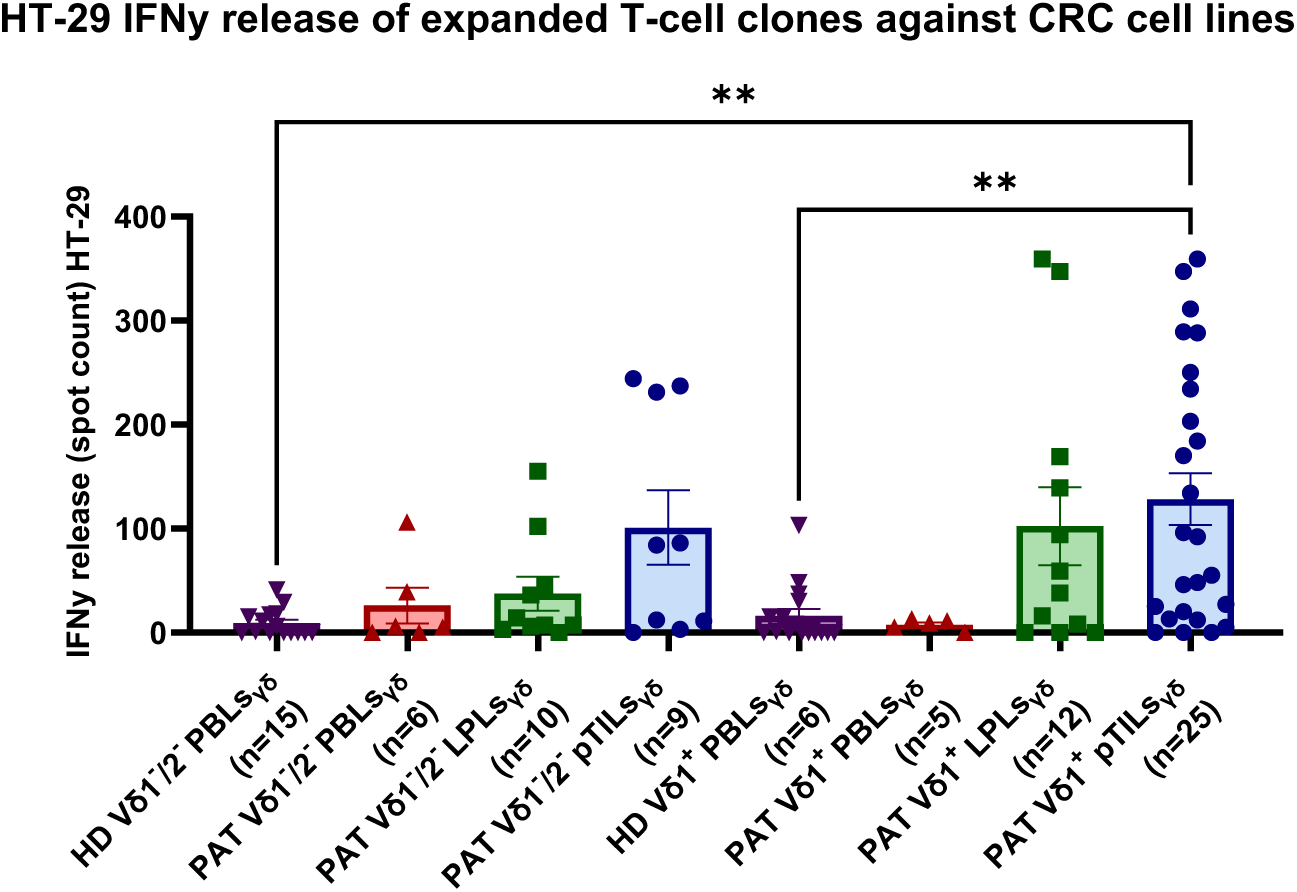
Tumor reactivity of single cell clones was determined by the production of 30 IFNγ spots after a 24-hour co-culture of T cells with one of the CRC cell lines at a 1:3 ratio, normalized to T cell only conditions. Statistical significance was determined by one-way ANOVA.

**Supplementary Fig. 10.**
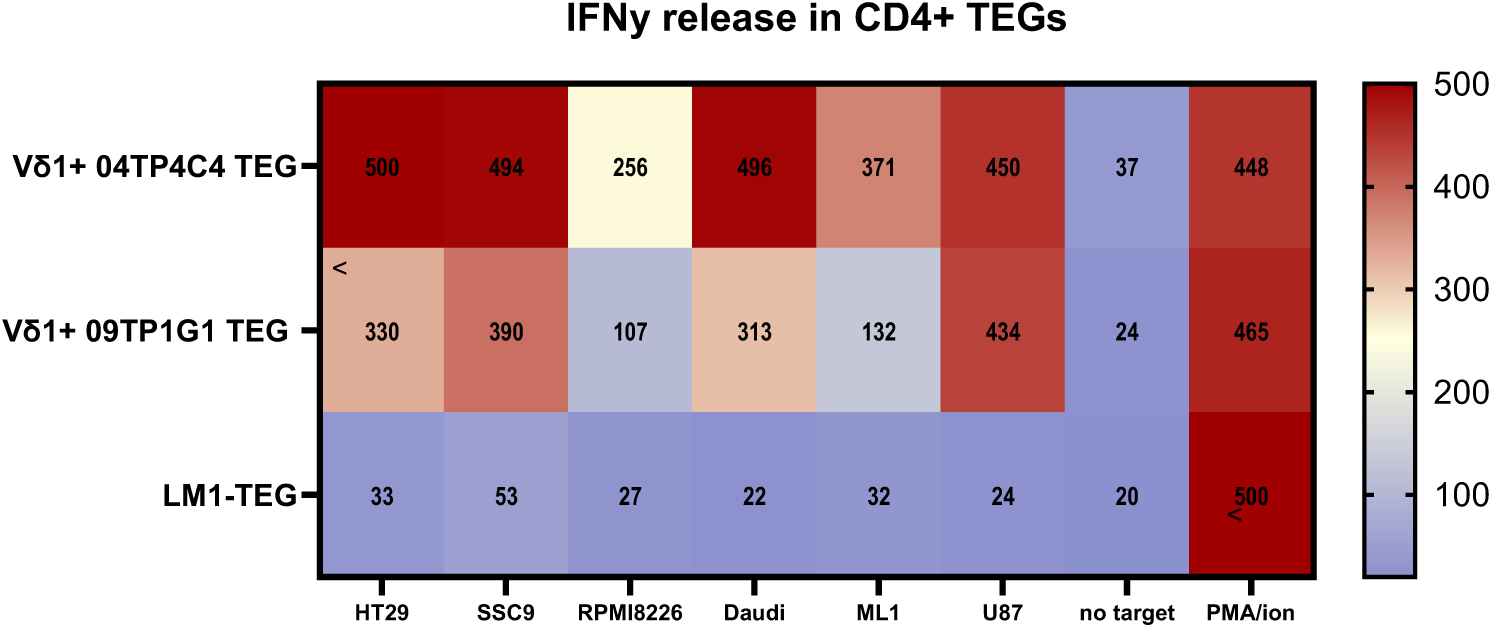
IFNγ release of tumor-reactive, Vδ1^+^ V09TP1G1 and 04TP4C4 TEGs compared to the mock TEG-LM1 upon a co-culture with a range of solid and hematological tumor cell lines, or PMA/Ionomycin as a positive control. Tumor reactivity was assessed after a 24-hour co-culture of T cells with the tumor cell lines at a 1:3 ratio, using an IFNγ ELISpot assay, and normalized to a no target control.

**Supplementary Fig. 11.**
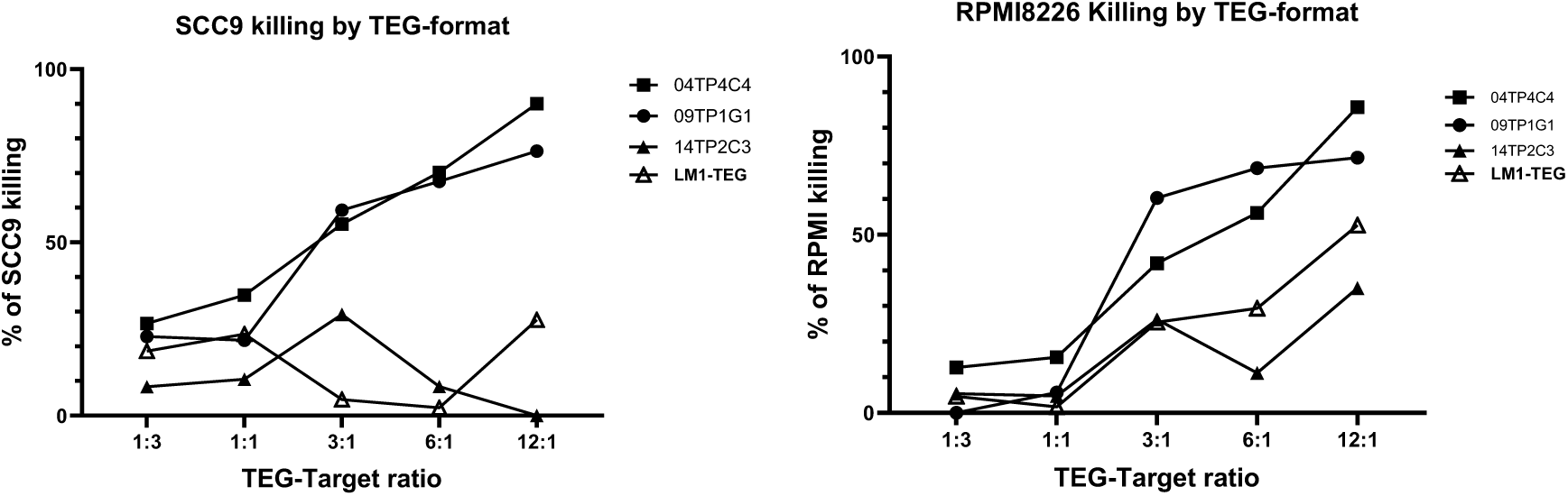
Luciferase based killing by the Vδ1+ 09TP1G1 and 04TP4C4 TEGs and the mock Vd2+ TEG (TEG-LM1) in CD4:CD8 1:1 ratio targeting the solid tumor cell line SCC9 and the hematological cell line RPMI8226 at multiple E:T ratios for 24h, normalized to the target only condition.

**Supplementary Fig. 12.**
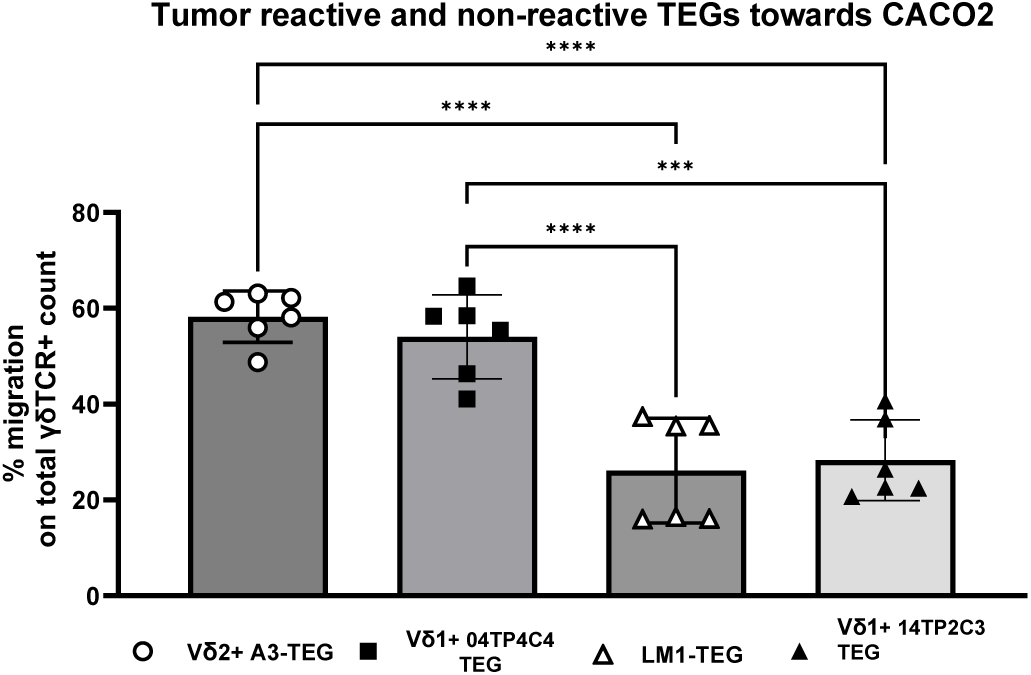
Migration dependent on tumor-reactive TCR. Percentage of migrated T cells of total T cells isolated at day 2 post T cell administration of either the tumor reactive Vδ2^+^ A3-TEG, Vδ1^+^ 04TP4C4 TEGs or non-reactive Vδ2^+^ TEG-LM1, Vδ1^+^ 14TP2C3 TEGs.

